# mRNA localization and thylakoid protein biogenesis in the filamentous heterocyst-forming cyanobacterium *Anabaena* sp. PCC 7120

**DOI:** 10.1101/2024.04.17.589878

**Authors:** Kexin Wang, Moontaha Mahbub, Giulia Mastroianni, Conrad W. Mullineaux

## Abstract

Heterocyst-forming cyanobacteria such as *Anabaena* (*Nostoc*) sp. PCC 7120 exhibit extensive remodeling of their thylakoid membranes during heterocyst differentiation. Here we investigate the sites of translation of thylakoid membrane proteins in *Anabaena* vegetative cells and developing heterocysts, using mRNA fluorescent *in situ* hybridization to detect the location of specific mRNA species. We probed mRNAs encoding reaction center core components and the heterocyst-specific terminal oxidases Cox2 and Cox3. As in unicellular cyanobacteria, we find that the mRNAs encoding membrane-integral thylakoid proteins are concentrated in patches at the inner face of the thylakoid membrane system, adjacent to the central cytoplasm. These patches mark the putative sites of translation and membrane insertion of these proteins. Oxidase activity in mature heterocysts is concentrated in the specialized “honeycomb” regions of the thylakoid membranes close to the cell poles. However, *cox2* and *cox3* mRNAs remain evenly distributed over the inner face of the thylakoids, implying that oxidase proteins migrate extensively after translation to reach their destination in the honeycomb membranes. The RNA binding protein RbpG is the closest *Anabaena* homolog of Rbp3 in the unicellular cyanobacterium *Synechocystis* sp. PCC 6803, which we previously showed to be crucial for the correct location of photosynthetic mRNAs. An *rbpG* null mutant shows decreased cellular levels of photosynthetic mRNAs and photosynthetic complexes, coupled with perturbations to thylakoid membrane organization and lower efficiency of the Photosystem II repair cycle. This suggests that chaperoning of photosynthetic mRNAs by RbpG is important for the correct co-ordination of thylakoid protein translation and assembly.

**IMPORTANCE:** Cyanobacteria have a complex thylakoid membrane system which is the site of the photosynthetic light reactions as well as most of the respiratory activity in the cell. Protein targeting to the thylakoids and the spatial organization of thylakoid protein biogenesis remain poorly understood. Some filamentous cyanobacteria show additional levels of complexity, with the differentiation of heterocysts, specialized nitrogen fixing cells in which the thylakoid membranes undergo extensive remodeling. Here we probe mRNA locations to reveal thylakoid translation sites in a heterocyst-forming cyanobacterium. We identify an RNA-binding protein important for the correct co-ordination of thylakoid protein translation and assembly, and we demonstrate the effectiveness of mRNA Fluorescent *in situ* Hybridization as a way to probe cell-specific gene expression in multicellular cyanobacteria.

## INTRODUCTION

*Anabaena* (or *Nostoc*) sp. PCC 7120 (hereafter *Anabaena*) is a multicellular, non-branching filamentous cyanobacterium that develops heterocysts in a semi-regular pattern along the filaments when it is not provided with combined nitrogen in the environment. Heterocysts are cells specialized for nitrogen fixation. They provide nitrogen compounds to vegetative cells and receive products of photosynthesis in return through cell-cell communication via septal junction complexes (1). The development of heterocysts from vegetative cells is conducted via a sophisticated regulatory network (2, 3). To provide a micro-oxic environment for nitrogen fixation in heterocysts, Photosystem II (PSII) is inactivated to halt photosynthetic oxygen production (4). The thylakoid membranes in developing heterocysts are re-organized to form two domains, the peripheral thylakoids (P domain) and the honeycomb thylakoids (H domain) with structural and compositional heterogeneity. The honeycomb membranes are clustered at the sub-polar regions of the cell near the cell junctions (5, 6) and are required to maintain the micro-oxic environment in the heterocyst cytoplasm by consuming oxygen that diffuses into heterocysts from the neighboring vegetative cells. This is achieved by the heterocyst-specific terminal oxidases Cox2 and Cox3 (Qox) (7, 8). By contrast, Cox1 is expressed specifically in vegetative cells as the sole terminal oxidase (9).

Most studies have focused on the differences in photosynthetic electron transport between heterocysts and vegetative cells, with the functionally distinct honeycomb thylakoids much less studied (5). Little is known about the process and molecular regulation of thylakoid membrane re-organization during heterocyst development. FraH, which is expressed at a later stage of heterocyst development, is suggested to be required for re-organization of intracellular membranes. FraH-GFP was observed at the polar region of heterocysts while expression remains in vegetative cells as well (10, 11). In a *fraH* null mutant, no honeycomb structure was developed and the nitrogenase activity was impaired almost completely in oxic environments (11). The role of *fraH* in the regulation of gene expression is unknown, but it is suggested that *fraH* is associated with the formation and maintenance of the honeycomb thylakoids (11). The overexpression of *fraH* was found to affect the organization of thylakoid membranes in vegetative cells, resulting in condensed membrane fragments (12). Several honeycomb-localized proteins are known, including HetN (13), Alr4119 (one of the two CurT homologs which are heterocyst honeycomb specific) (14) and Valyl-tRNA synthetase (15). Specific sequences involved in the targeting of proteins were further studied. Residues 2 to 27 from the N-terminal of HetN were found to be responsible for the localization of the protein at honeycomb thylakoids (13). In mature heterocysts, Valyl-tRNA synthetase (ValRS) is anchored to the membrane by a C-terminal CAAD domain and located specifically in the honeycomb membranes (14).

Recent work on the unicellular cyanobacterium *Synechocystis* sp PCC 6803 (hereafter *Synechocystis*) has implicated RNA binding proteins (RBPs) in delivering mRNAs encoding thylakoid proteins to the thylakoid surface, with possible roles in targeting proteins to the thylakoids rather than the plasma membrane (16). In heterocyst-forming cyanobacteria such as *Anabaena*, the development of honeycomb membranes in heterocysts adds additional complexity to the protein targeting problem. Here, we use mRNA Fluorescent *in situ* Hybridization (FISH) (17) to probe the location of several thylakoid membrane mRNAs in *Anabaena*. We demonstrate that FISH provides an excellent new approach to probing cell-specific gene expression in differentiating filaments. We show that mRNAs encoding thylakoid proteins are clustered at the proximal surfaces of the thylakoid system. The mRNAs encoding the honeycomb membrane proteins Cox2 and Cox3 are similarly distributed to the mRNAs encoding photosystem subunits: they do not show preferential location near the poles of the cell. This implicates protein migration in formation of the honeycomb membranes, rather than targeting prior to translation. In *Synechocystis*, the RNA binding protein Rbp3 is implicated in binding photosynthetic mRNAs and their location at the thylakoid membranes (16). Here we show that an *Anabaena* knockout mutant lacking RbpG, the nearest *Anabaena* homolog of Rbp3, shows highly perturbed thylakoid membrane organization and function, underlining the role of RBPs in thylakoid biogenesis.

## RESULTS

### Photosynthetic mRNA localization

RNA FISH was shown in single-celled cyanobacteria to be effective as a probe for mRNA expression and localization, although the background fluorescence in cyanobacteria causes problems with the detection of scarce mRNAs (16). We used the *Anabaena* genome sequence (18) to design FISH probes against two mRNA species encoding membrane-integral thylakoid proteins: *psaA* (alr5154) and *psbA1* (alr4866), encoding core subunits of Photosystem I (PSI) and PSII respectively. We designed sets of 47-48 oligonucleotide probes against the coding sequences, as specified in Tables S1 and S2. *Anabaena* has 5 *psbA* genes expressed to different extents and in different circumstances (19). We designed our probes against the *psbA1* gene, which is often the most expressed (19). However, the gene family shows strong sequence conservation at the nucleotide level, so our FISH probes will certainly recognize other *psbA* mRNAs when present, as demonstrated in *Synechocystis* (16). The FISH probes were labelled with 5-Carboxytetramethylrhodamine (TAMRA) which provides a stable fluorescence signature distinct from that of the photosynthetic pigments (16).

In undifferentiated *Anabaena* filaments grown in nitrate-containing medium, *psaA* and *psbA* mRNAs were observed in most cells and localized in clusters at the inner surfaces of the thylakoid membrane system that face the central cytoplasm (Fig 1A). This is consistent with the pattern seen in the unicellular cyanobacteria *Synechocystis* and *Synechococcus* (16). As in the unicellular cyanobacteria, it suggests that these proximal thylakoid surfaces may be the main site of translation of photosynthetic proteins. To test this idea, we examined the effect of puromycin, an antibiotic that blocks translation and releases ribosomes from the mRNA (20). Puromycin treatment results in greatly increased cellular levels of *psbA* mRNA and a more diffuse distribution of this mRNA species (Fig 1B). The strong perturbation caused by puromycin implies that the FISH signals seen in untreated cells highlight mRNA that is coupled to ribosomes engaged in active translation. As in unicellular cyanobacteria (16), *psbA* mRNA remains at least partially located close to the proximal thylakoid surfaces in the presence of puromycin (Fig 1B), which suggests that there is a ribosome-independent factor that targets this mRNA to the thylakoid surface.

**FIG 1.**
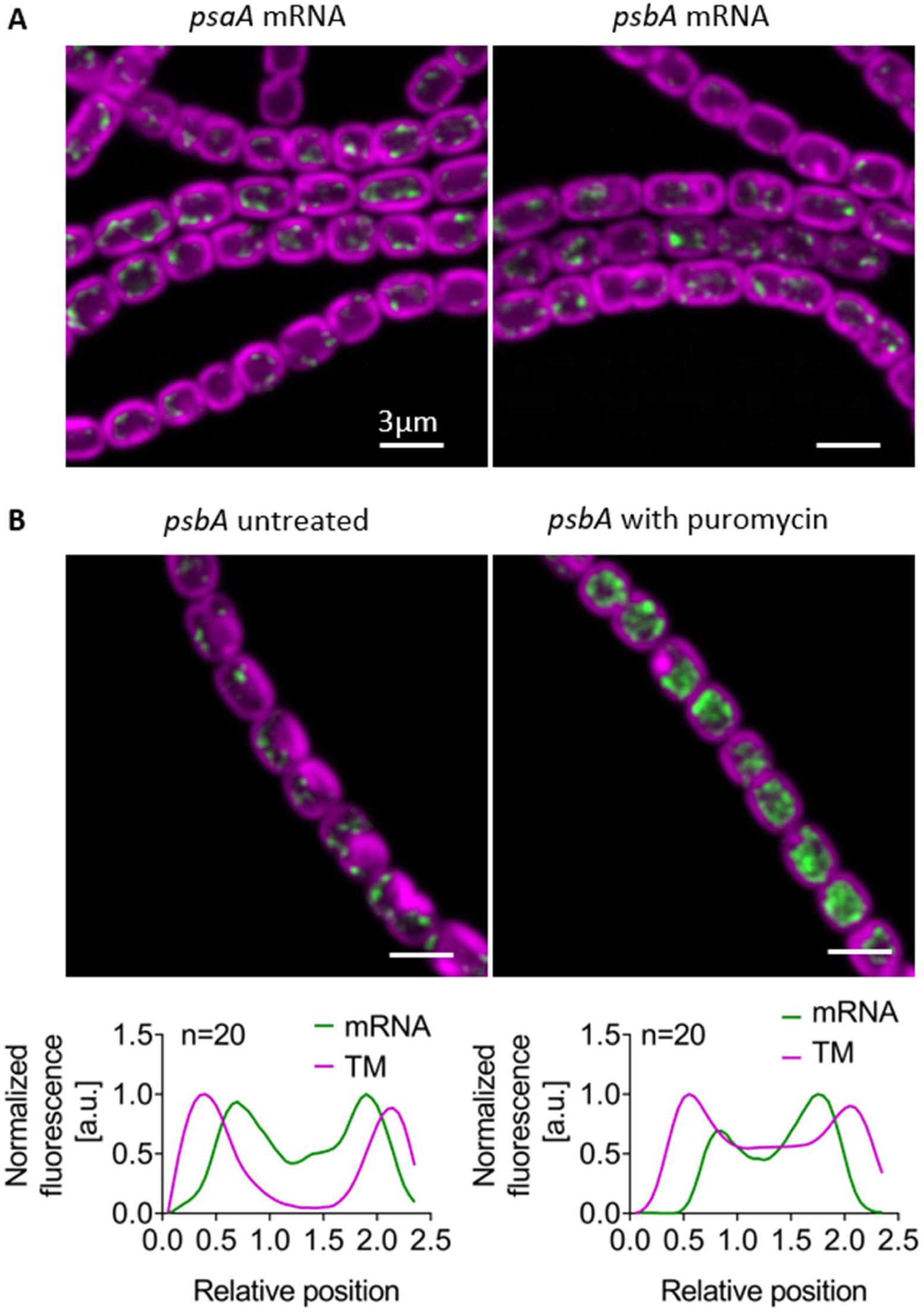
Localization of photosynthetic mRNAs in *Anabaena*. Confocal images showing TAMRA-labelled FISH probes for *psaA* and *psbA* in green, photosynthetic pigments in the thylakoids (TM) in magenta. Cells were grown in BG11 medium. Scale-bars: 3µm. **A**. Comparison of *psaA* and *psbA* signals; **B**. Effect of puromycin on the *psbA* signal, with averaged line profiles across 20 cells shown below.

### Heterocyst differentiation

To examine cell-specific gene expression and mRNA location in differentiating *Anabaena* filaments, we probed *psbA* mRNA as a function of time in filaments transferred to nitrate-free BG11_0_ medium to induce heterocyst formation (21). In addition to *psbA*, we probed the specifically heterocyst-expressed *cox2* and *cox3* gene clusters, encoding thylakoid membrane oxidases (9). Cox2 is a standard *aa*_3_-type cytochrome *c* oxidase (9, 22) whereas Cox3 is a *bo*_3_-type quinol oxidase categorized as a type-1 Alternative Respiratory Terminal Oxidase (ARTO) and recently referred to as Qox (22). Sets of 48 TAMRA-labelled probes were designed against the first 1500 coding bases of *coxB2A2* mRNA (comprising the alr2514 ORF and the first part of alr2515) and *coxB3A3* mRNA (comprising the alr2731 ORF and the first part of alr2732) (Tables S4, S5). We tested the specificity of the FISH probes with *cox2* and *cox3* null mutants (9). In strain CSAV140 (*coxB2*::C.S3) a segment of the *coxB2* gene is replaced with an antibiotic resistance cassette (9). As expected, no *cox2* FISH signal was detectable in this mutant (Fig S1A). By contrast, in strain CSAV135 (*coxA3*::C.S3) (9) we unexpectedly found a heterocyst-specific *cox3* FISH signal at comparable levels to the wild type (Fig S1B). However, the insertional site in this mutant is downstream of the region probed, and so this may simply indicate that the upstream mRNA remains sufficiently expressed and stable for detection.

Up to 6 hours after nitrogen stepdown by transfer to BG11_0_ medium, *psbA* FISH signals were detectable in all cells and no *cox2* expression was detectable (Fig 2). However, after 8 hours some cells lacked *psbA* signals and strong *cox2* expression could be observed in some cells (Fig 2). The *cox2*-expressing cells were always well spaced in the filament and generally showed lower photosynthetic pigment fluorescence, indicating likely pro-heterocysts. The *cox2* gene cluster was previously reported to be induced by about 9h after nitrogen deprivation and peak by about 12h under the regulation of NtcA, a global nitrogen regulator (9). We found that we could reliably probe mRNA levels in developing heterocysts up to ∼20 h after nitrogen stepdown, but were never able to obtain heterocyst FISH signals at 24 h (eg Fig 3B). This is most likely because the developing multi-layered heterocyst envelope (21) prevents effective permeabilization of the cells. Unexpectedly, however, we found the FISH probing was again possible at 48 h (Fig 3B).

**FIG 2.**
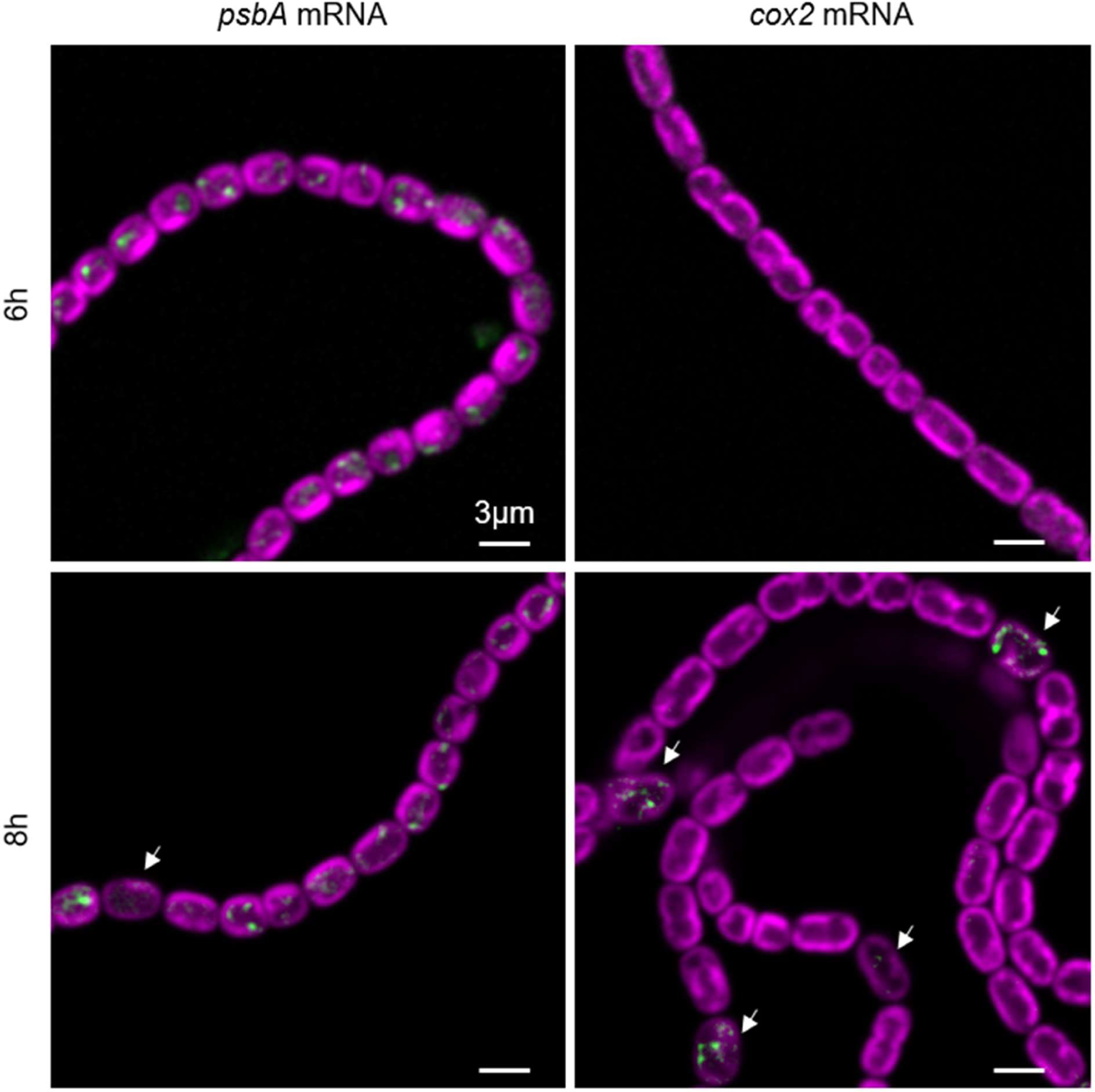
Effect of nitrogen step-down on *psbA* and *cox2* mRNAs. Confocal images showing TAMRA-labelled FISH probes for *psbA* in green, photosynthetic pigments in the thylakoids in magenta. Cells were fixed at 6 h or 8 h after transfer to BG11_0_ medium. Arrows indicate likely pro-heterocysts. Scale-bars: 3µm

**FIG 3.**
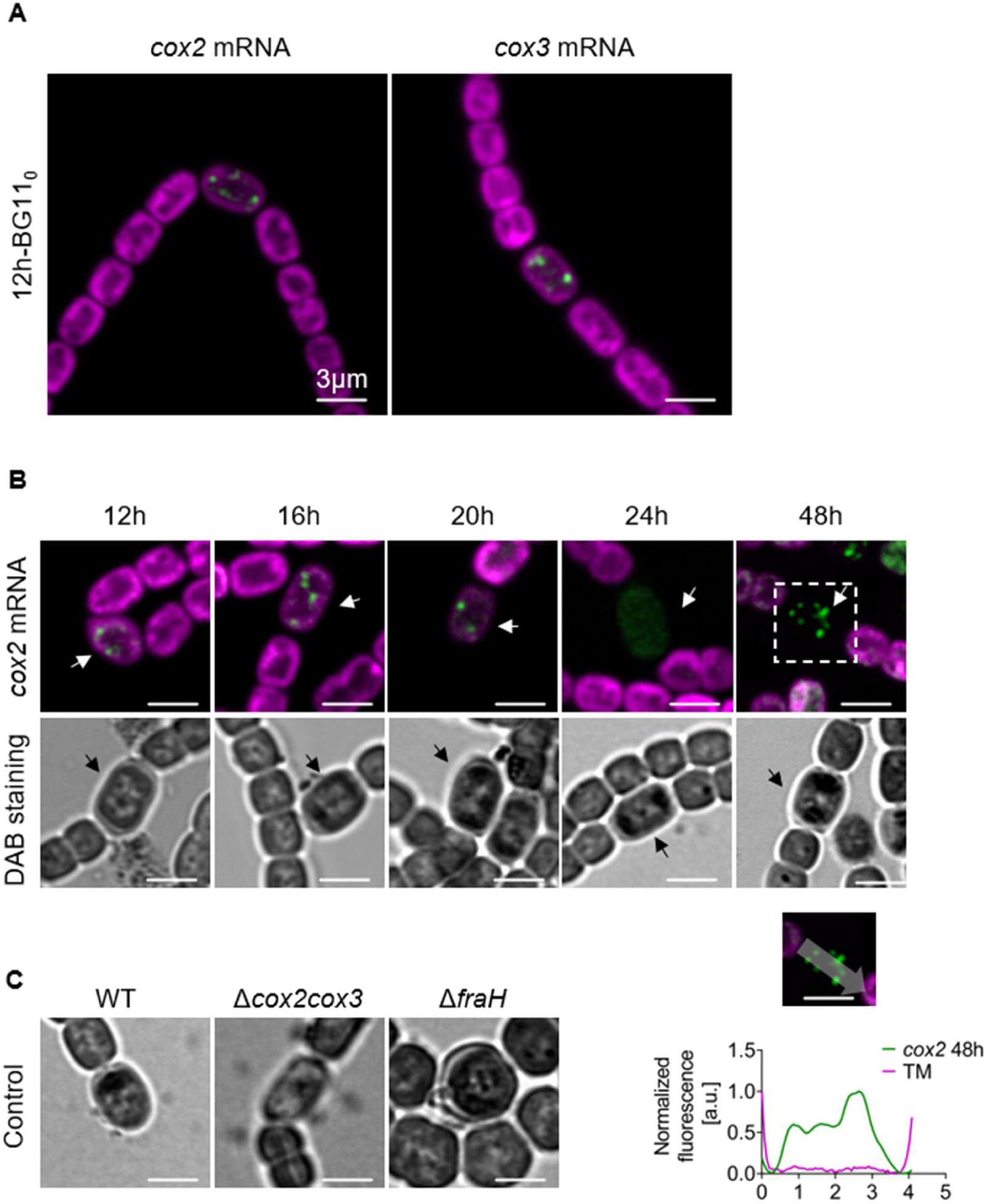
Localization of *cox* mRNAs and oxidase activity in developing *Anabaena* heterocysts. Confocal images showing TAMRA-labelled FISH probes for *cox2* and *cox3* in green, photosynthetic pigments in the thylakoids in magenta. Arrows indicate pro-heterocysts or heterocysts. **A**. *cox2* and *cox3* mRNA localization 12h after transfer to BG11_0_ medium. **B**. Location of *cox2* mRNA (top panel) and oxidase activity from DAB staining (bottom panel) at different times after transfer to BG11_0_ medium. A line profile of *cox2* mRNA signal along the long axis of a heterocyst is shown below. **C.** Oxidase activity revealed by DAB staining in heterocysts of wild type *Anabaena* and the Δ*cox2cox3* and Δ*fraH* mutants. Cells were collected after 24 h in BG11_0_ medium. Scale-bars: 3µm.

### Thylakoid remodeling in heterocysts

We probed the location of *cox2* mRNA as a function of time after nitrogen stepdown (Fig 3). At all timepoints from 6-48 h, *cox2* mRNA showed a patchy location at the proximal edge of the thylakoids (Figs 2,3), similar to the location of *psaA* and *psbA* mRNAs in vegetative cells (Figs 1,2). *cox3* mRNA showed a similar distribution at the timepoints we tested (Fig 3A, Fig S1B). Even in fully mature heterocysts at 48 h, there was no discernible bias in *cox2* mRNA localization towards the cell poles (Fig 3B). In parallel with the FISH measurements, we probed the location of oxidase activity in developing heterocysts with diaminobenzidine (DAB) staining (7). DAB is oxidized to form a brown precipitate which can be visualized by brightfield microscopy (7). In vegetative cells and young pro-heterocysts, oxidase activity was evenly distributed around the cell periphery, but in maturing heterocysts it was increasingly concentrated in the sub-polar regions (Fig 3B), reflecting the formation of the honeycomb membranes (7). To check that Cox2 and Cox3 oxidases were responsible for the DAB staining that we observed in heterocysts, we tested the CSAV 141 mutant lacking both oxidases (9) and saw no comparable DAB signal in heterocysts (Fig 3C). We also tested a Δ*fraH* mutant which does not develop honeycomb membranes (11), and saw oxidase activity only at the peripheral thylakoids (Fig 3C).

### RNA binding proteins in *Anabaena*

In *Synechocystis*, Rbp2 and Rbp3, two members of a family of RNA binding proteins, were shown to bind photosynthetic mRNAs and implicated in targeting to the thylakoid membranes (16). *Anabaena* has eight members of a homologous family of RBPs, encoded by the *rbpA1*, *rbpA2*, *rbpB*, *rbpC*, *rbpD*, *rbpE*, *rbpF* and *rbpG* genes (23). All these RBPs contain a conserved RNA recognition motif in the N-terminal region but are more divergent at the C-terminus. One subgroup, termed Class 1 Type 1, have a glycine-rich C-terminal extension, including Rbp1 in *Synechocystis* and RbpA1, RbpA2, RbpB, RbpC, RbpE, and RbpF in *Anabaena* (24, 25). *Synechocystis* Rbp2 and *Anabaena* RbpD belong to Class I type II, which lack the glycine-rich region. *Synechocystis* Rbp3 and *Anabaena* RbpG both have a longer C-terminal extension that is not rich in glycine, both have been assigned to Class II (24–26). We carried out a multiple alignment of amino acid sequences of the 8 *Anabaena* RBPs and the 3 *Synechocystis* RBPs (Fig S2A) and constructed an evolutionary phylogenetic tree based on the aligned sequences (Fig S2B). The results confirm the close homology of *Synechocystis* Rbp3 and *Anabaena* RbpG.

### Phenotype of *Anabaena* Δ*rbpG*

We constructed an *rbpG* null mutant, in which the coding sequence was replaced by a neomycin resistance cassette. The mutant was fully segregated (Fig S3) after 4 months’ growth on neomycin, with a final concentration at 200 µg/ml. Full segregation of an *rbpG* null mutant was previously reported not to be achievable (23), and it is possible that our Δ*rbpG* strain may have acquired a compensating secondary mutation to allow complete loss of *rbpG*. Because of the homology between *Anabaena* RbpG and *Synechocystis* Rbp3, which is implicated in photosystem biogenesis (16), we probed the photosynthetic phenotype of Δ*rbpG*.

Confocal imaging of phycocyanin (PC) and chlorophyll (Chl) fluorescence for cells grown in standard BG11 medium show striking changes in thylakoid membrane organization (Fig 4A). Wild type cells show a thick and rather irregular layer of thylakoid membranes around the cell periphery, often with loops of membrane extending into the central cytoplasm. In Δ*rbpG* the peripheral layer of thylakoids appears thinner and more regular, but many cells show in addition very bright fluorescent spots showing strong fluorescence from PC as well as Chl (Fig 4A). On average, Δ*rbpG* cells showed lower Chl and PC fluorescence (Fig 4B), and absorption spectra indicate a lower cellular pigment content (Fig 4C). 77K fluorescence spectra with chlorophyll excitation (Fig 4D) show the same shape in Δ*rbpG* as in wild type, indicating no major change in the PSII/PSI ratio and suggesting no major perturbation in reaction center assembly. 77K spectra with phycobilin excitation, however, show significantly higher fluorescence from PC and allophycocyanin relative to the PSII and PSI reaction centers (Fig 4D), despite the lower PC/Chl ratio in the cells (Fig 4C). This implies some perturbation in energy transfer from the phycobilisomes to the reaction centers. The Δ*rbpG* mutation also affects the dimensions of the cells, with Δ*rbpG* cells bring significantly shorter and wider than those of the wild type (Fig 4E). To probe the effects of the Δ*rbpG* mutation on cell architecture at higher resolution, we examined cells by super-resolution fluorescence microscopy and thin-section electron microscopy (Fig 5). Super-resolution imaging of fluorescence from the photosynthetic pigments (lattice SIM^2^ with resolution of about 60 nm in the *xy* plane) showed that the concentrated spots of pigment fluorescence were linked to the rest of the thylakoid membranes, either to the peripheral thylakoids (as in Cell 1, Fig 5A) or to the membranes in the sub-polar region (as in Cell 2, Fig 5A). Thin-section electron micrographs (Fig 5B) confirm the thinner peripheral thylakoid zone in Δ*rbpG*, with significantly fewer thylakoid membrane layers on average (Fig 5C). Δ*rbpG* cells generally lack the moderately curved thylakoid zones that can occupy much of the cytoplasm in wild type cells, but sometimes show localized highly curved protrusions of the thylakoids into the central cytoplasm (examples shown in Fig 5B). These most likely correspond to the highly fluorescent patches seen in Δ*rbpG* (Figs 4A, 5A).

**FIG 4.**
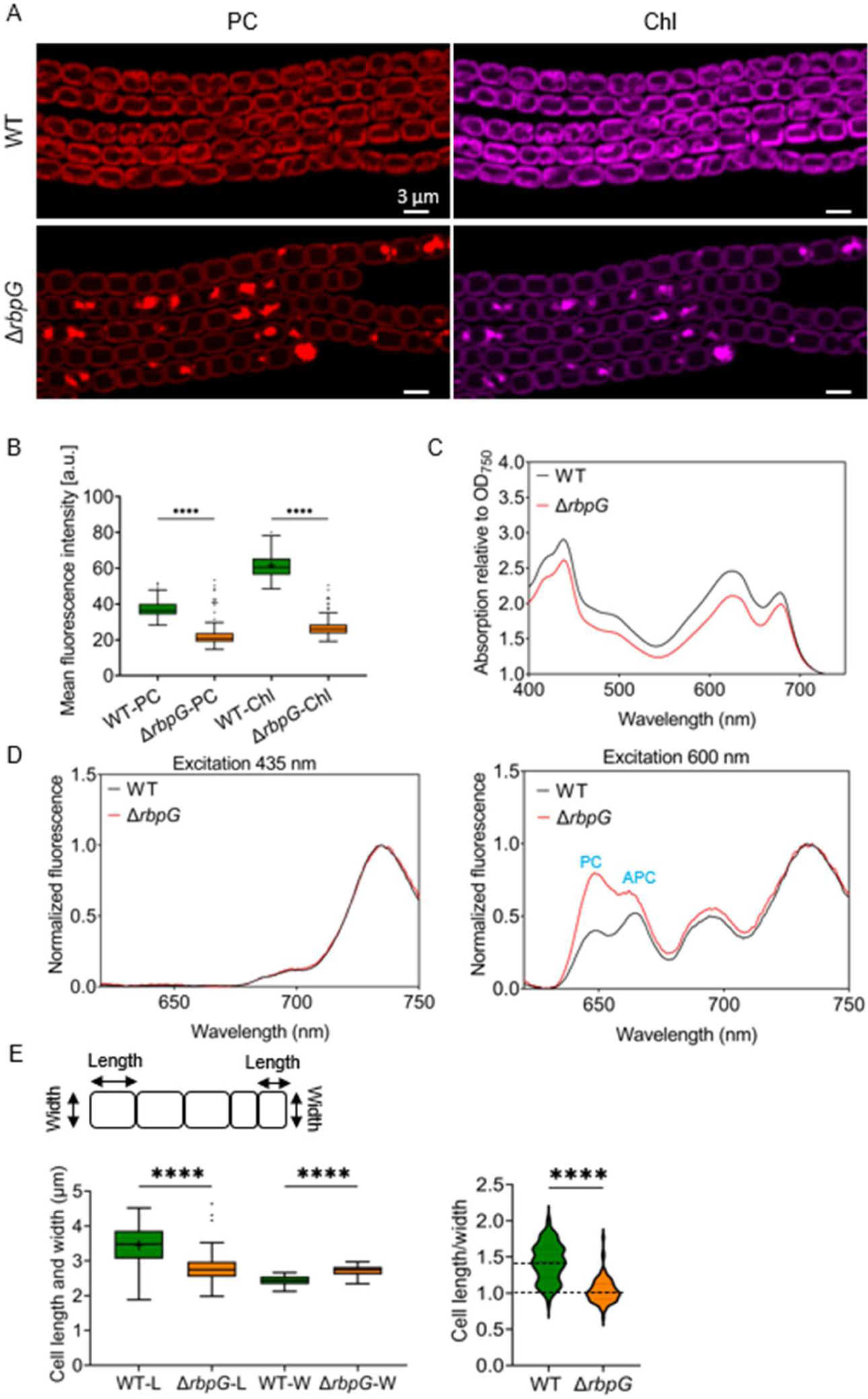
Phenotype of the Δ*rbpG* mutant. **A.** Confocal images of phycocyanin (PC) and chlorophyll (Chl) fluorescence. Scale-bars: 3 µm. **B.** Mean cellular PC and Chl fluorescence intensities in the wild type and the Δ*rbpG* mutant (n=100 cells). **C.** Absorption spectra normalized to turbidity at 750 nm (averaged from 3 biological replicates). **D.** 77K fluorescence emission spectra at 77K with 435 nm (Chl) and 600 nm (PC) excitation, normalized to the Photosystem I peak (averaged from 3 biological replicates). **E.** Cell dimensions: length (L), width (W) and L:W ratio (n=100 cells). **** indicates statistical significance (p < 0.0001).

**FIG 5.**
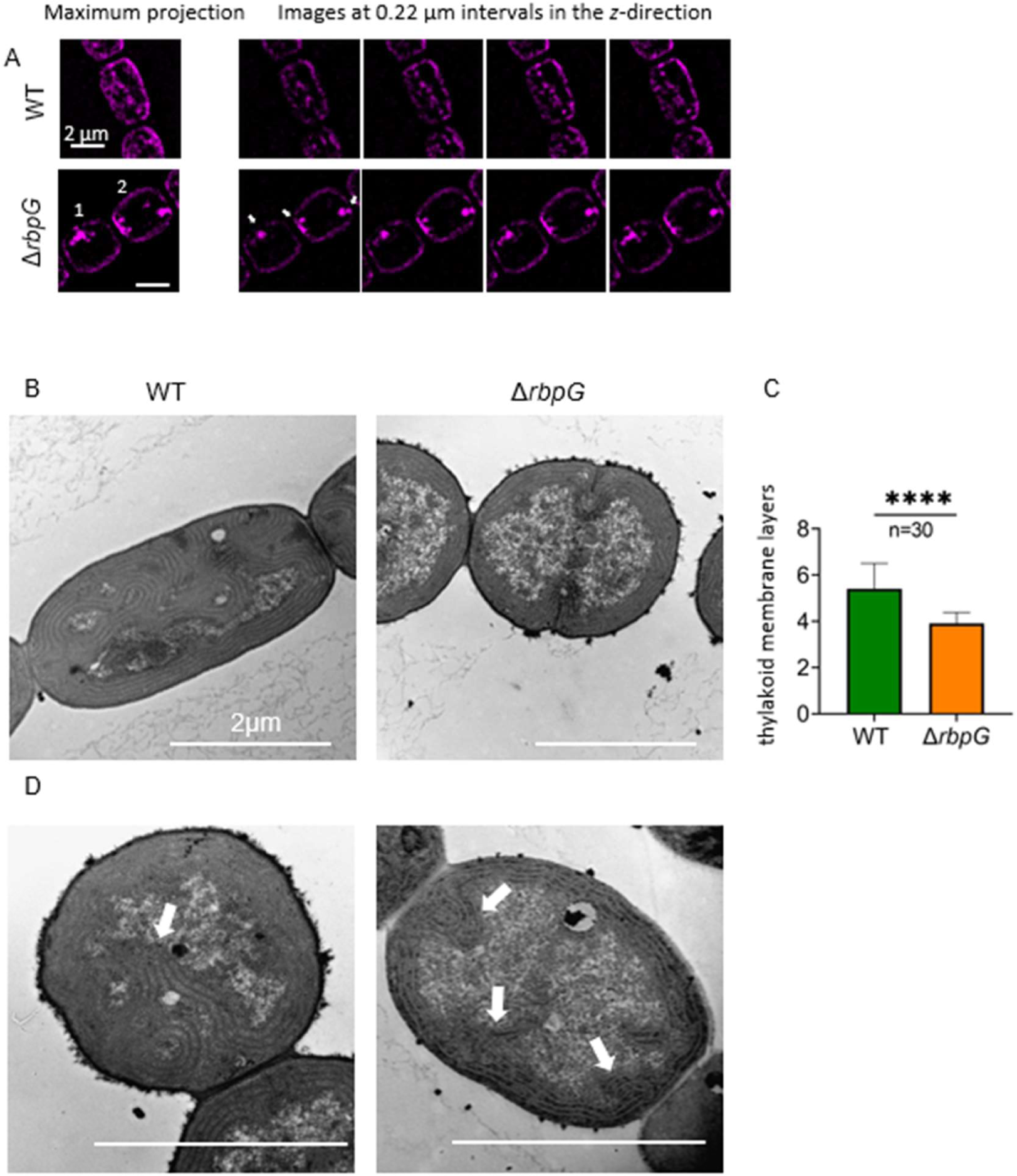
Thylakoid membrane conformation of the Δ*rbpG* mutant at higher resolution. **A** Super-resolution (SIM^2^) imaging of chlorophyll fluorescence in *Anabaena* wild type and Δ*rbpG* cells. The arrows highlight dense spots of chlorophyll fluorescence. Scale-bars: 2 µm. **B.** Representative thin-section electron micrographs of *Anabaena* wild type and Δ*rbpG* cells. **C.** Quantitation of number of thylakoid membrane layers from thin-section electron micrographs (30 cells of each strain). **** indicates statistical significance (p < 0.0001). **D.** Representative examples of thin-section electron micrographs of Δ*rbpG* cells that show highly curved and condensed thylakoid membrane regions that may correspond to the bright patches seen in fluorescence micrographs (arrows highlight these regions). Scale-bars: 2 µm.

### mRNA levels and the PS II repair cycle in Δ*rbpG*

We use FISH to probe the levels and location of photosynthetic mRNAs in the Δ*rbpG* mutant. *psbA* and *psaA* mRNAs were probed as before (Fig 1) and in addition we probed the *cpcAB* transcript encoding the α- and β-phycocyanin subunits of the phycobilisomes, using a further set of TAMRA-labeled probes (Table S3). The mean cellular FISH signals from all 3 of these probes were significantly decreased in Δ*rbpG* as compared to wild type, with a particularly strong decrease in *psbA* signal (Fig 6 A,B). However, the sub-cellular distribution of the mRNAs was not obviously affected, with most FISH foci being located at the proximal edge of the thylakoid system in both wild type and Δ*rbpG* (Fig 6A). We also probed for *cox2* mRNA as previously (Fig 3) but saw no clear differences between Δ*rbpG* and wild type (Fig 6C). *cox2* FISH signals remained heterocyst-specific and with similar levels and location to the wild type (Fig 6C).

**FIG 6.**
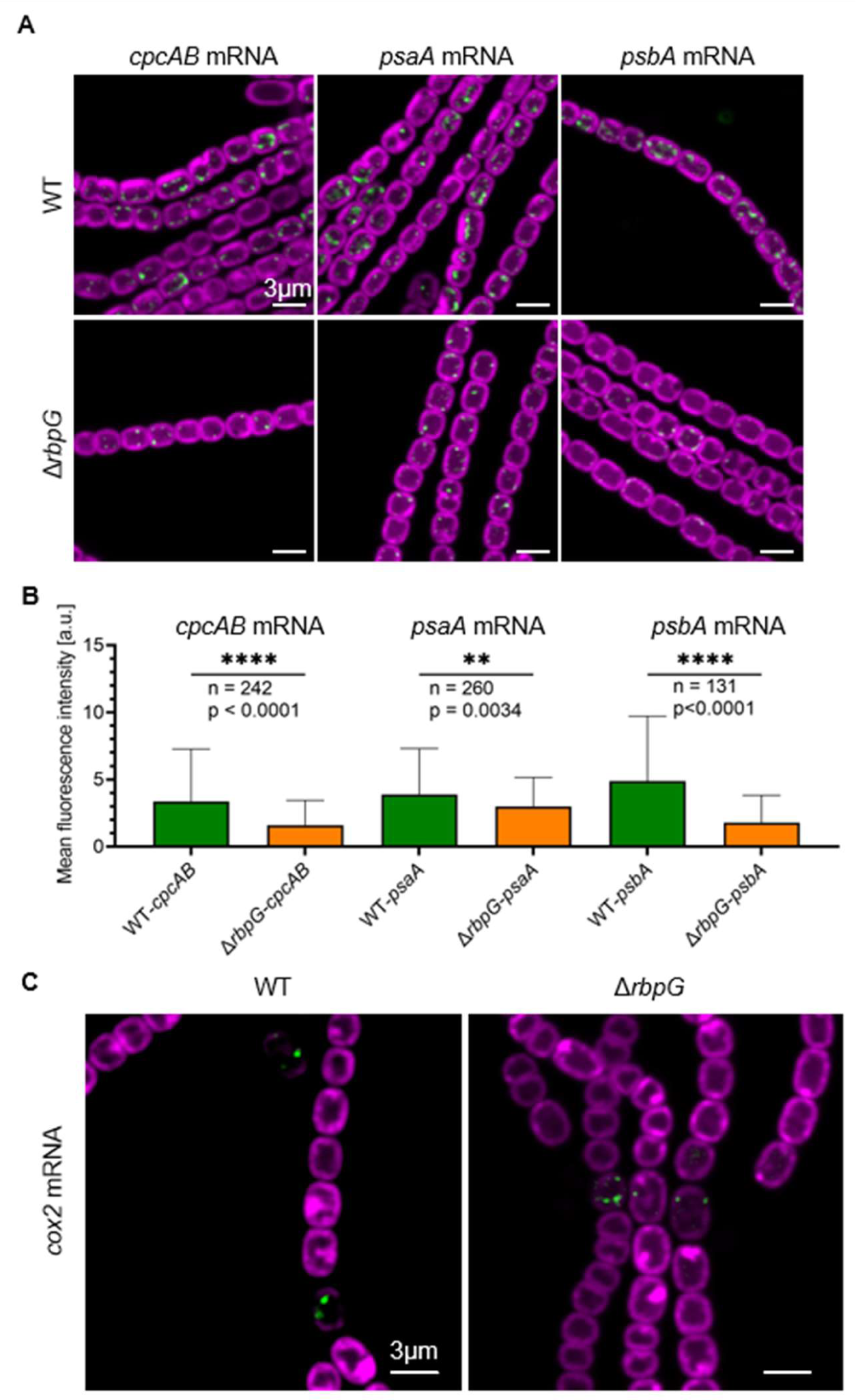
Effects of the Δ*rbpG* mutation on mRNA localization and expression. TAMRA FISH signals are shown in green and fluorescence from the photosynthetic pigments in magenta. All scale-bars: 3 µm **A.** Localization and expression of 3 photosynthetic mRNAs in Δ*rbpG* vs. wild type *Anabaena* (cells grown in standard BG11 medium). **B**. Mean cellular fluorescence intensity of FISH signals for the 3 photosynthetic mRNAs in wild type vs. Δ*rbpG. n =* number of cells measured; **** = significant difference with p < 0.0001; ** = significant difference with p < 0.01. **C.** *cox2* mRNA FISH signals in wild type *Anabaena* and Δ*rbpG* (cells fixed 16h after transfer to BG11_0_ medium).

Given the pronounced effects of the Δ*rbpG* mutation on *psbA* mRNA levels (Fig 6 A,B) and thylakoid membrane conformation (Fig 4A, Fig 5) we tested for effects of the Δ*rbpG* mutation on the PSII repair cycle, which requires rapid synthesis and integration of new PsbA subunits to replace those affected by photodamage (27). We tested for PS II oxygen-evolving activity in an oxygen electrode using saturating light and an excess of the artificial PS II electron acceptor 2,6-dichloro-1,4-benzoquinone (DCBQ) (28). Δ*rbpG* cells grown in moderate light showed lower PSII activity (per Chl) than wild type (Fig 7). Exposure to high light for 20 min resulted in significant decreases in PSII activity, to a similar extent in wild type and Δ*rbpG*. However, after transfer back to moderate light the wild type recovered PSII activity almost completely within 40 min, whereas Δ*rbpG* recovered only about 40% of the lost activity in this time. This indicates that the efficiency of the PSII repair cycle is lower in the Δ*rbpG* mutant.

**FIG 7.**
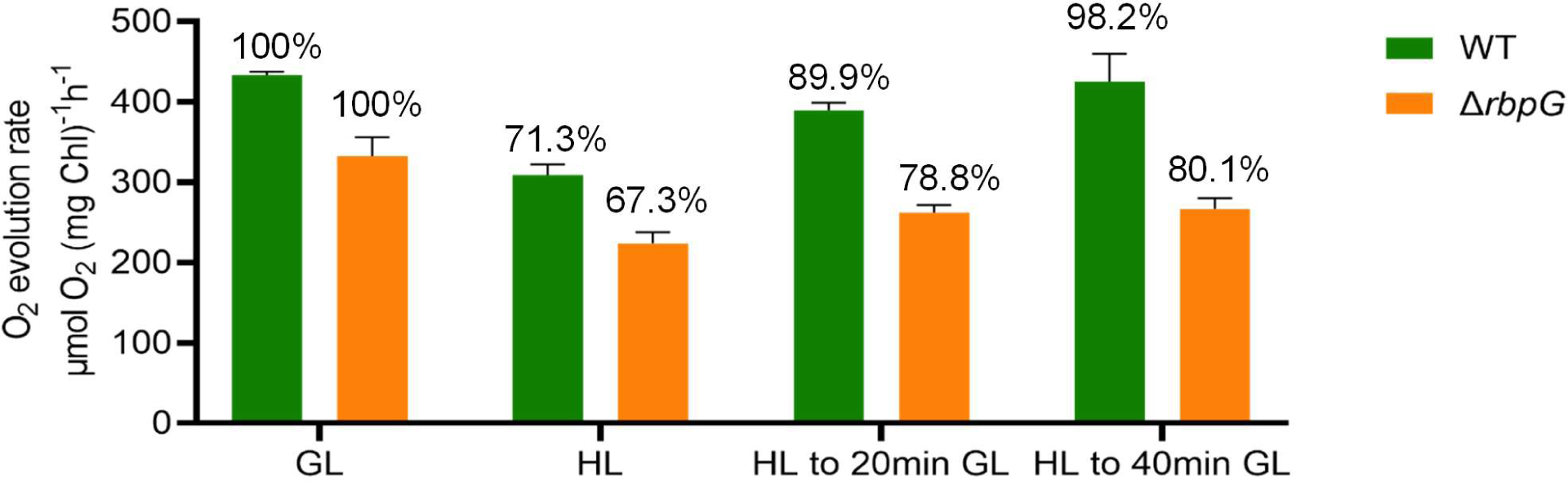
PSII activity and efficiency of the PSII repair cycle in *Anabaena* wild type and Δ*rbpG*. O_2_ evolution was measured at saturating light in the presence of excess PSII electron acceptors. Cells were exposed to high light (HL) for 20 min, then returned to growth light (GL). The bars indicate rates of O_2_ evolution per chlorophyll, and the percentages above indicate rates relative to the initial rate before exposure to HL. All values are means from 3 technical replicates, with SD indicates by the error bars.

### Δ*rbpF* mutant and Δ*rbpF/G* double mutant

*Anabaena* RbpF is an RBP belonging to Cluster 1 with homology to *Synechocystis* Rbp1 (Fig S2). Transcriptomic analysis indicates that its expression is under the control of the heterocyst regulator HetR, and therefore it is likely to be specifically expressed in heterocysts (29). We constructed a Δ*rbpF* null mutant and a Δ*rbpF/G* double mutant. Both mutants were fully segregated (Fig S3). We looked for effects of the mutations on *psbA*, *psaA* and *cox2* mRNAs but could not detect any effects of the Δ*rbpF* mutation. The Δ*rbpF* mutant resembled the wild type, while Δ*rbpF/G* resembled Δ*rbpG* (Fig S4).

## DISCUSSION

### RNA-FISH as a probe for cell-specific gene expression in multicellular cyanobacteria

Filamentous cyanobacteria show multiple kinds of cell differentiation, of which heterocyst development is the most studied. Cell-specific gene expression has generally been probed by the use of GFP or luciferase reporter genes fused to promoters of interest (eg (30–32)). Here we show that RNA-FISH can provide a more direct readout of cell-specific mRNA levels: the technique confirms, for example, the loss of *psbA* expression and the induction of *cox2* and *cox3* expression in developing heterocysts (Fig 2). The use of FISH has some limitations, however. As discussed in (16), transcripts should ideally be at least ∼1000 bases long. The high background autofluorescence in cyanobacteria precludes the detection of low-abundance mRNAs and can cause difficulties with quantitation of the signal. In unicellular cyanobacteria, we found that background fluorescence in the channel used for TAMRA can reliably be predicted from the fluorescence signal in the red channel used for the photosynthetic pigments, allowing background fluorescence to be subtracted out from the FISH signal (16). This technique also works well in vegetative cells of *Anabaena*. However, we found that the fluorescence ratio in the two channels is different in heterocysts, and changes unpredictably during heterocyst development. Finally, the thick heterocyst cell envelope seems to be prevent permeabilization for access of the FISH probes beyond ∼20 h after nitrogen stepdown, although, unexpectedly, FISH signals can again be detected in older heterocysts at 48 h (Fig 3). This suggests greater permeability of the envelope in older heterocysts. A population of apparently ageing heterocysts showing reduced metabolite exchange with their vegetative cell neighbors is detectable 48 h after nitrogen stepdown (1). Our FISH results suggest that ageing heterocysts may also lose some integrity in their cell envelope, however gene expression is still active.

### Sites of thylakoid protein translation in *Anabaena*

We previously examined the locations of mRNA species encoding photosynthetic proteins in two species of unicellular cyanobacteria, *Synechocystis* and *Synechococcus elongatus* PCC 7942 (hereafter *Synechococcus*). In both cases, we found photosynthetic mRNAs clustered at the innermost surface of the thylakoid system (16). Photosynthetic mRNA species in *Anabaena* are similarly distributed (Figs 1,2) although the cell shape is distinct from both the spherical *Synechocystis* and the thin, rod-shaped *Synechococcus*. The pattern is not confined to photosystem mRNAs, or to vegetative cells, as mRNAs encoding the heterocyst-specific thylakoid membrane-located oxidases Cox2 and Cox3 show a similar location (Fig 2, Fig 3). This suggests that the proximal thylakoid surface is a universal site of thylakoid protein translation in cyanobacteria. In agreement with the idea that mRNA located there is being actively translated, we found that treatment with the translation inhibitor puromycin changes the distribution of *psbA* mRNA, which becomes more abundant and more widely spread over the thylakoid surface (Fig 1B). Most mRNAs are destabilized by puromycin treatment, but *psbA* mRNA abundance consistently increases with puromycin treatment, as previously seen in *Synechocystis* and *Synechococcus* (16). This hints at a highly-conserved mechanism of regulation of PsbA biogenesis based on the promotion of *psbA* mRNA degradation by translation (33, 34). As in *Synechocystis* and *Synechococcus* (16), *Anabaena psbA* mRNA retains some affinity for the thylakoid surface even in the presence of puromycin (Fig 1B), implying a translation-independent mode of targeting of the mRNA to the thylakoid. Such translation-independent targeting of mRNAs encoding membrane proteins has also been observed in *E. coli* (35).

### Remodeling of thylakoid membranes in heterocysts

Thylakoid membranes undergo extensive remodeling during heterocyst development, with segregation into the peripheral thylakoids and the sub-polar honeycomb membranes (6). These membranes have distinct protein compositions, but they must be physically continuous as a Fluorescence Recovery after Photobleaching (FRAP) measurement showed that GFP-tagged AtpA can diffuse from pole to pole in mature heterocysts (6). We confirmed that terminal oxidase activity in heterocysts is strongly concentrated in the honeycomb region (Fig 3B), as previously observed (7). The polar concentrations of oxidase activity are not observed in a *cox2cox3* null mutant (Fig 3C) and therefore they must reflect the location of active Cox2 and Cox3 oxidases. Santamaría-Gómez *et al* proposed complementary mechanisms by which proteins could accumulate specifically at the honeycomb membranes: by diffusion from the peripheral thylakoids and capture by a trapping factor, or by specific protein insertion in the honeycomb membrane (6). Migration of entire membranes also seems to contribute to formation of the honeycomb membrane structure (6). In the case of Cox2 and Cox3, we find no evidence for translation specifically at the honeycomb membranes, since the mRNAs are located in foci that remain distributed across the entire proximal surface of the thylakoid system at all stages of heterocyst development, with no detectable bias to the polar regions even in mature heterocysts with well-established honeycomb membranes (Fig 3). Our results suggest that Cox2 and Cox3 proteins are translated at multiple points across the inner surface of the thylakoid system and accumulate at the honeycomb membranes by diffusion and capture. Our recent results in *Synechococcus* suggest that membrane proteins must be freely mobile in the biogenic proximal thylakoid surface where mRNAs and ribosomes are located, since different PS II subunits are translated at different locations and therefore must diffuse to find their interaction partners (36).

### Role of RbpG in thylakoid membrane biogenesis

*Anabaena* RbpG is a member of a conserved family of cyanobacterial mRNA binding proteins with strong homology to *Synechocystis* Rbp3, which was shown to bind photosynthetic mRNAs and is implicated in the correct location of *psbA* and *psaA* mRNAs at the thylakoid surface (16). Our *Anabaena ΔrbpG* knockout shows a range of phenotypic features consistent with a role in promoting the efficient translation of photosynthetic mRNAs. Levels of *cpcAB*, *psaA* and *psbA* mRNAs are all significantly reduced in *ΔrbpG*, with the strongest effect on *psbA* mRNA levels (Fig 6). In low light-grown cells, the levels of photosynthetic pigments are lower in *ΔrbpG* than in wild type (Fig 4), and there is a marked reduction in the efficiency of the PSII repair cycle (Fig 7), which restores PSII activity after photodamage in high light and which requires rapid biogenesis and integration of new PsbA subunits (27). The mutant shows striking changes in thylakoid membrane conformation (Figs 4,5). The distribution of photosynthetic pigment-protein complexes in the membranes is often very uneven in *ΔrbpG*, with strong accumulation in occasional highly-curved protrusions of the membranes (Figs 4,5). The simplest explanation for the multiple phenotypic effects of *ΔrbpG* is that the RbpG protein binds a set of photosynthetic mRNAs, as does the homologous Rbp3 in *Synechocystis* (16). RbpG could then chaperone those mRNA species, preventing their premature degradation and thereby increasing the efficiency of thylakoid protein biogenesis. The changes in thylakoid ultrastructure and photosynthetic complex distribution in *ΔrbpG* further suggest a role in the spatial coordination of biogenesis of the photosynthetic machinery. In *ΔrbpG*, there appears to be a frequent overaccumulation of photosystems in specific confined and highly curved regions of the thylakoid. A link between membrane curvature (induced by the CurT protein) and PSII biogenesis has been noted in *Synechocystis* (37). Our results suggest that mRNA chaperoning by *Anabaena* RbpG helps to distribute photosystem biogenesis across the thylakoid surface, preventing overaccumulation in highly localized membrane areas. It remains to be determined whether other members of the *Anabaena* RBP family have related functions. Here, we knocked out *RbpF* but saw no comparable phenotypic effects (Fig S4).

## MATERIALS AND METHODS

### Cyanobacterial cultivation

Cells were cultivated in BG11 medium (38) supplemented with TES-NaOH buffer (pH 8.2). NaHCO_3_ was added at a final concentration of 10mM. Liquid cultures were maintained in tissue culture flasks (Sarstedt) at 30 °C under constant white light (5-10 µmol photons m^−2^ s^−1^) at 100 r.p.m. Cyanobacterial cultures were also maintained on BG11 plates with 1.5% Bacto Agar (BD Difco™). Antibiotics were added for the cultivation of some mutants when indicated at the following concentrations: neomycin, 10 µg ml^−1^; spectinomycin, 2 µg ml^−1^; streptomycin, 2 µg ml^−1^. Cyanobacterial strains used in this study are shown in Table 1. To induce heterocyst formation, *Anabaena* cells were harvested by centrifugation (2500 × g) and washed three times with 800µl BG11_0_ medium (38). The cells were then maintained in BG11_0_ using conical flasks to avoid breakage of filaments with heterocysts.

**Table 1.**
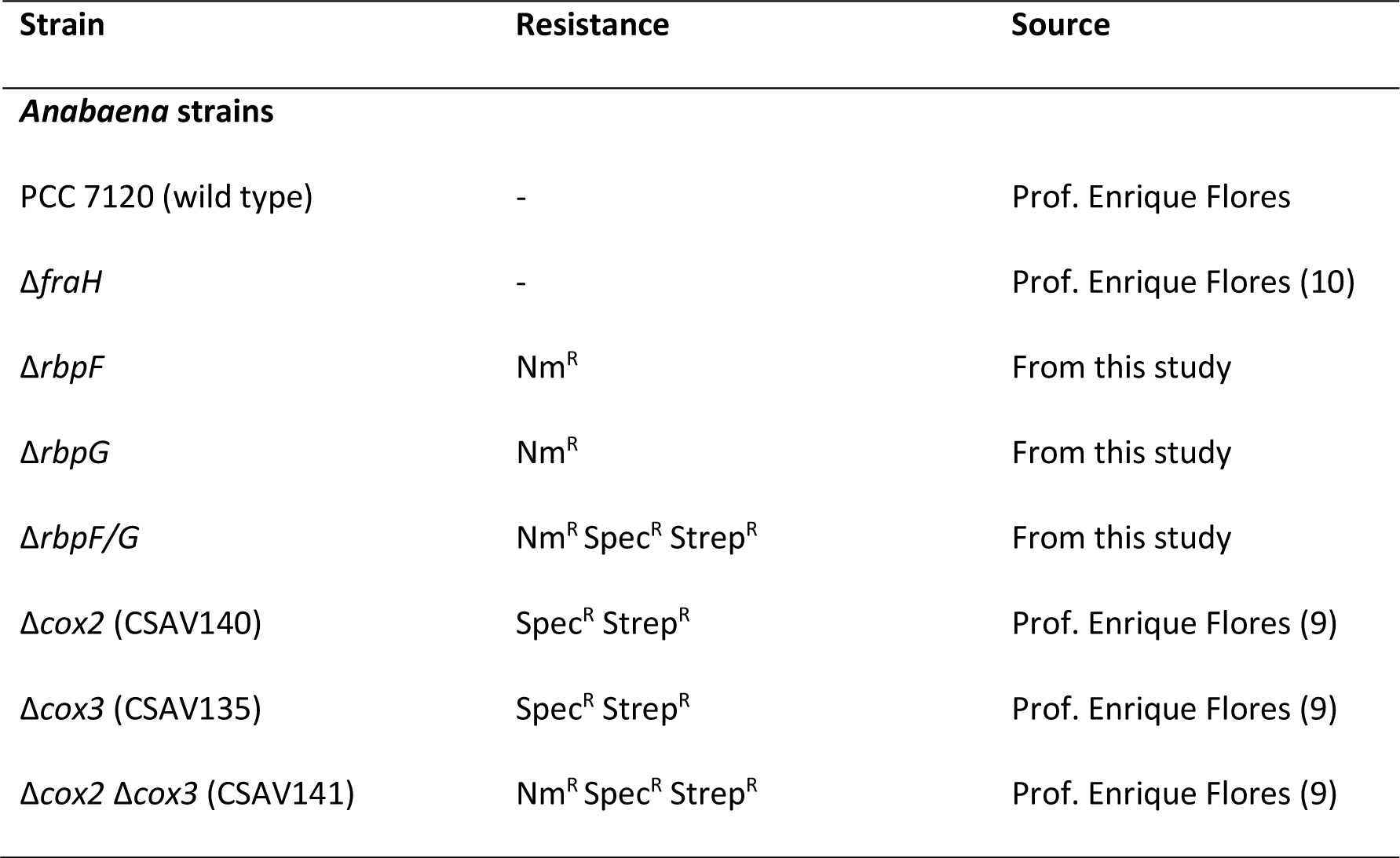
Cyanobacterial strains used in this study.

### Design of FISH probes

FISH probes were designed against the target mRNAs by using Stellaris® Probe Designer version 4.2 (Biosearch Technologies, https://www.biosearchtech.com/stellaris-designer). The coding sequence of target gene was obtained from the complete genome of *Nostoc* sp. PCC 7120 available at NCBI (National Center for Biotechnology Information, https://www.ncbi.nlm.nih.gov/nuccore/BA000019.2 (18). We designed sets of 40-48 TAMRA-labeled probes, each 20 nucleotides long, ∼50% GC content and with a minimum spacing of 2 bases between hybridization sites. Masking Level 2 was used to avoid general problematic RNA sequences. The probe sets were purchased from LGC Biosearch Technologies. Probes for *psaA*, *psbAI*, *cpcAB*, *cox2*, *cox3* mRNAs are listed in Figs S1-S5.

### mRNA-FISH (RNA-fluorescent *in situ* hybridization)

Samples were prepared using an adapted version of the original protocol by Skinner *et al* (17). The protocol was as previously described in (16) except that the permeabilization step was extended: formaldehyde-fixed cells were permeabilized in 70% ethanol for 1 h on a nutator at room temperature, followed by 3-5 h at 4 °C and then an additional hour at room temperature. *Anabaena* cells were collected after cultivation in liquid BG11 or BG11_0_ medium. For puromycin treatment, cultures were incubated for 1 h at 30 °C in 1ml liquid medium supplemented with puromycin at a final concentration of 500 µg/ml, followed by formaldehyde fixation.

### Mutagenesis

Deletion mutants were generated in which the gene of interest was replaced with an antibiotic-resistant cassette by double homologous recombination. A neomycin/kanamycin-resistance cassette (Nm^R^) was used to replace *rbpG* (all4377) in the Δ*rbpG* mutant. 1000 bp sequences upstream and downstream of the *rbpG* coding sequence were cloned by colony PCR. The amplified upstream, Nm^R^ and downstream sequences were integrated by Gibson assembly into a version of the backbone vector pRL271B modified with BamHI and PstI restriction sites. The constructed cargo plasmid was introduced into *E.coli* donor HB101 cells containing a helper plasmid pRL623 (39) and then introduced into *Anabaena* cells by conjugation (40) with a second *E. coli* strain ED8654 containing the conjugative plasmid pRL443 (39). *Anabaena* cultures were sonicated to generate shorter filaments. Cell culture containing 10 µg Chl *a* was collected and mixed with pre-mixed *E. coli* cultures for conjugation. The *sacB* gene from the plasmid pRL271B was used as a marker for selection of double recombination. In Δ*rbpF*, the coding sequence plus 22 bp of downstream sequence was replaced by a neomycin/kanamycin-resistance cassette (Nm^R^). In Δ*rbpF/G*, the same sequence in the Δ*rbpG* background was replaced by a spectinomycin/streptomycin-resistance cassette (Spec^R^ Strep^R^). Full segregation of the mutants was checked by colony PCR (Fig S3). *E. coli* strains and plasmids are listed in Table S6 and PCR primers are listed in Table S7.

### Confocal microscopy and image processing

Samples were prepared by spotting 5-10 µl of cell suspension onto a 1.5% agarose plate (low melt agarose in 1× PBS). Images were obtained with a TCS-SP5 laser scanning confocal microscope using an HCX PL APO 63× oil immersion objective (numerical aperture 1.4). Excitation was provided by a 561 nm DPSS laser. Fluorescence emission was collected from 565-580 nm for TAMRA and 660-700 nm for photosynthetic pigments. The confocal pinhole was set to give an optical section in the *z*-direction of ∼0.72 µm, and images were acquired at a scan speed of 400 Hz with 16 × line averaging and a 1024 × 1024 pixel format with 12 bit depth. Image processing was with Fiji ImageJ (https://imagej.net/Fiji/Downloads (41).

### Super-resolution microscopy

Images were collected with a Zeiss Elyra 7 microscope with Structured Illumination Microscopy (Lattice SIM^2^). Imaging used a Plan-Apochromat 63×/1.4 oil DIC M27 objective and high definition 0.17mm thickness coverslips. Excitation was at 488 nm and detection at 570-620 nm. The exposure time was 50 ms and the bit depth was 16.

### Transmission electron microscopy

*Anabaena* cultures were collected and fixed for 2h with 4% glutaraldehyde in Sørensen’s phosphate buffer at pH 7.3. Cells were then postfixed with 1% osmium tetroxide and dehydrated with a series of increasing ethanol concentrations (50% to 100%), embedded in resin, and cut into thin sections of 70 nm. The thin sections were then post-stained with UA-Zero EM stain (Agar Scientific) for 10 min and Reynolds’ lead citrate for 4 min, and were washed with distilled water. Images were recorded using a JEOL model JEM-1400 Flash transmission electron microscope.

### Absorption and fluorescence spectroscopy

Absorption spectra were obtained using an SLM Aminco DW-2000 spectrophotometer. Cyanobacterial cultures were grown to mid-log phase. Spectra were recorded from 400 nm to 750 nm and were normalised to turbidity at 750 nm. Data presented are means of three biological replicates. Chlorophyll concentrations were estimated from the absorption spectra using an extinction coefficient of 68 mM^−1^cm^−1^ and deconvoluting according to (*A*Chl_678_ = 1.0162 × *A_67_*_8_ - 0.0630 × *A_625_*) (42).

Fluorescence emission spectra of cyanobacteria samples were recorded at 77K with a Perkin Elmer LS55 fluorescence spectrometer equipped with a liquid nitrogen housing. All samples were concentrated to 5 µM Chl in BG11 medium and loaded into silica capillary tubes. The tubes were dark-adapted for 5 min and then snap frozen in liquid nitrogen. Fluorescence emission spectra were recorded from 620-750 nm with slit-width 3-5 nm and excitation at 435 nm for Chl *a* and 600 nm for phycocyanin. Spectra were normalized to the PSI peak (∼735 nm) and were averaged from 3 biological replicates.

### DAB staining for oxidase activity

To visualize the activity of terminal oxidases in (pro-)heterocysts, cultures were collected by centrifugation at specific time points after transfer to BG11_0_ medium. The cells were mixed with freshly prepared 1× DAB (3,3′-diaminobenzidine, 0.5 mg/mL) using the DAB Substrate Kit (10x, Pierce™). The samples were then incubated in the dark for 1 h (7) before imaging with the Leica TCS-SP5 confocal microscope in brightfield mode using a 458 nm Argon laser line.

### Oxygen evolution measurements

Oxygen evolution was measured in Clarke-type oxygen electrode (OxyLab2, Hansatech, UK). Cyanobacterial cultures were concentrated to 10 µM Chl in BG11 medium and dark-adapted for 10 min before measurements. PSII activity was assessed from oxygen evolution at 30 °C in the presence of 2 mM 2,6-dichloro-1,4-benzoquinone (DCBQ) and 1 mM potassium ferricyanide (28) and saturating light at approximately 35000 μmol photons m^−2^ s^−1^ (Schott KL 2500 LED projector). Cultures were photoinhibited in high light (HL) (white light at 1000-1500 μmol photons m^−2^ s^−1^) at 30 °C for 20 min. Cells were then transferred back to growth light (GL) for the recovery phase. Data presented are means of 3 technical replicates.

### Phylogenetic tree reconstruction

Amino acid sequences of RNA-binding proteins in *Anabaena* sp. PCC 7120 and *Synechocystis* sp. PCC 6803 were retrieved from the Kyoto Encyclopedia of Genes and Genomes (KEGG) database. Multiple alignment was carried out using the ClustalW program (Clustal Multiple alignment – Graphic view) (43) on BioEdit version 7.2.6 (44). The phylogenetic tree was constructed using the Minimum Evolution method (45) in MEGA X (46) employing the Poisson correction model (47). The robustness of the phylogenetic tree was confirmed by bootstrap analysis based on 500 replicates.

## SUPPLEMENTARY MATERIAL

## ACKNOWLEDGMENTS

Research was supported by Biotechnology and Biological Sciences Research Council (UKRI BBSRC) research grants BB/W001012/1 and BB/T017716/1. KW was supported by a scholarship (#201904910467) from the China Scholarship Council. We thank Enrique Flores (IBVF, Sevilla) for the provision of wild type *Anabaena* and several mutants constructed in his lab, Dennis Nürnberg and Fabian Conradi (Freie Universität Berlin) for plasmids for *Anabaena* transformation, Isidora Zagradanin for FISH experiments with *cox3* probes and Annegret Wilde (University of Freiburg) for critical reading of the manuscript.

## SUPPLEMENTARY MATERIAL

**FIG S1.**
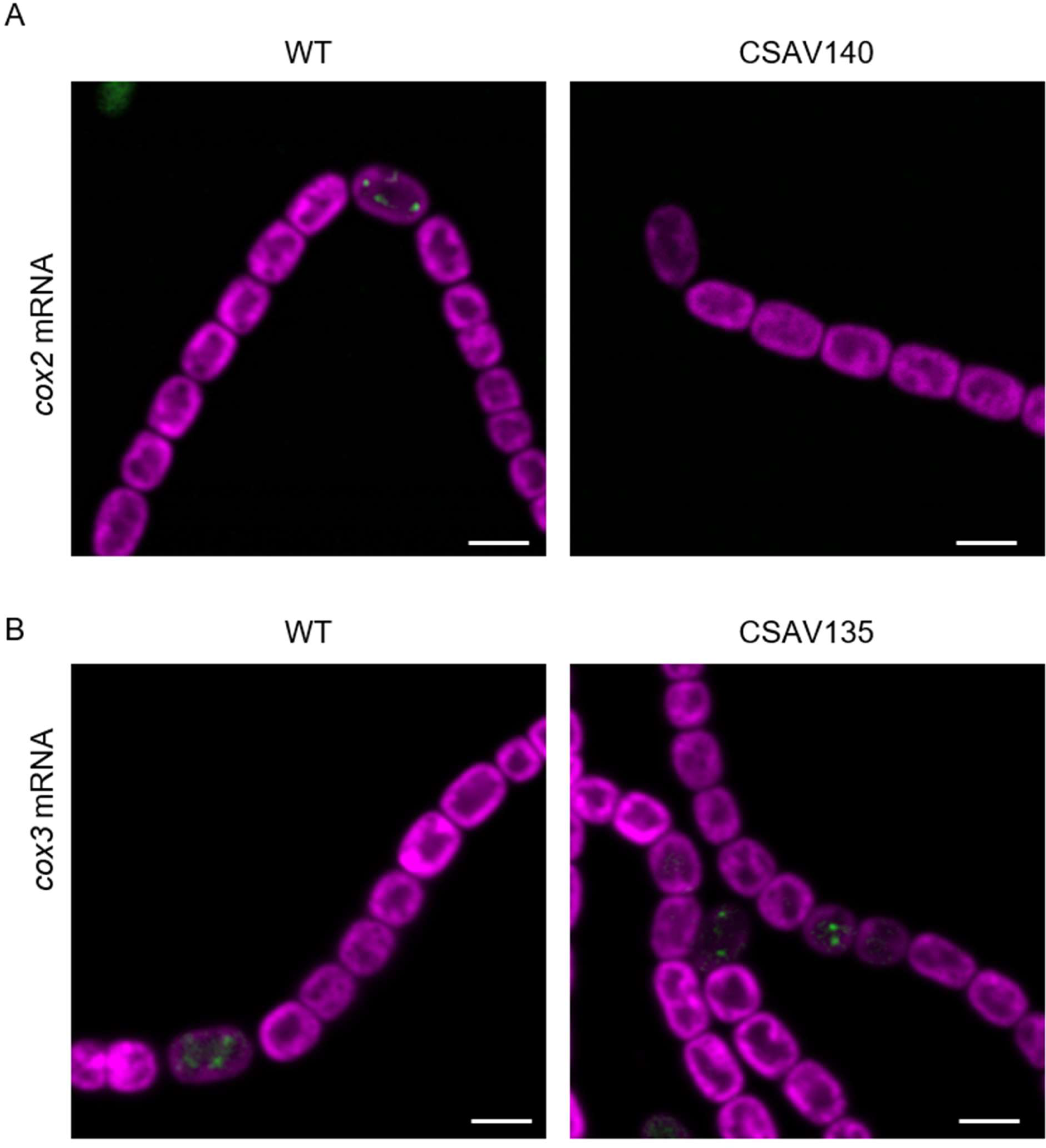
*cox* mRNA FISH signals in *cox* mutants.

**FIG S2.**
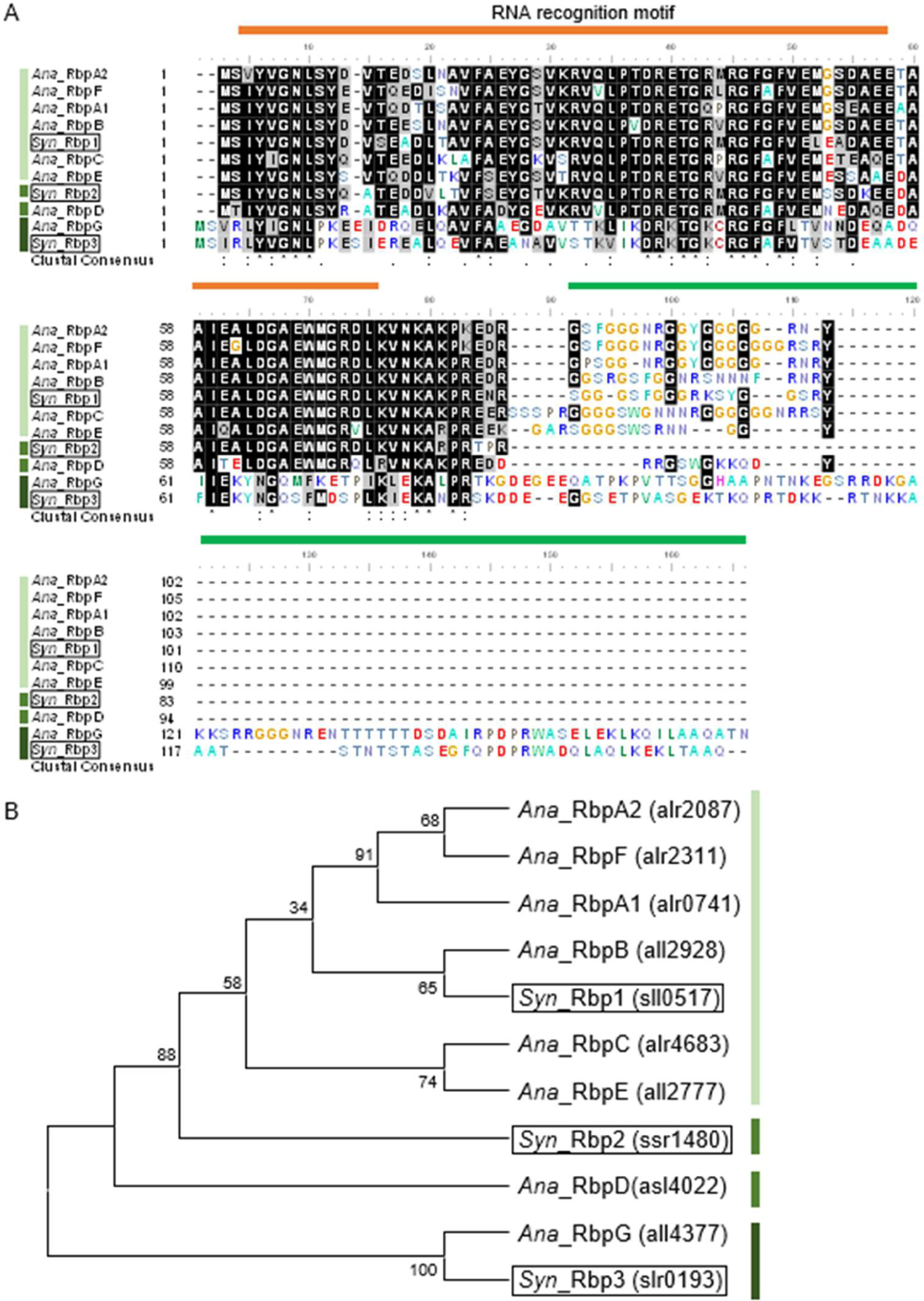
Phylogenetic tree of *Anabaena* and *Synechocystis* RBP sequences.

**FIG S3.**
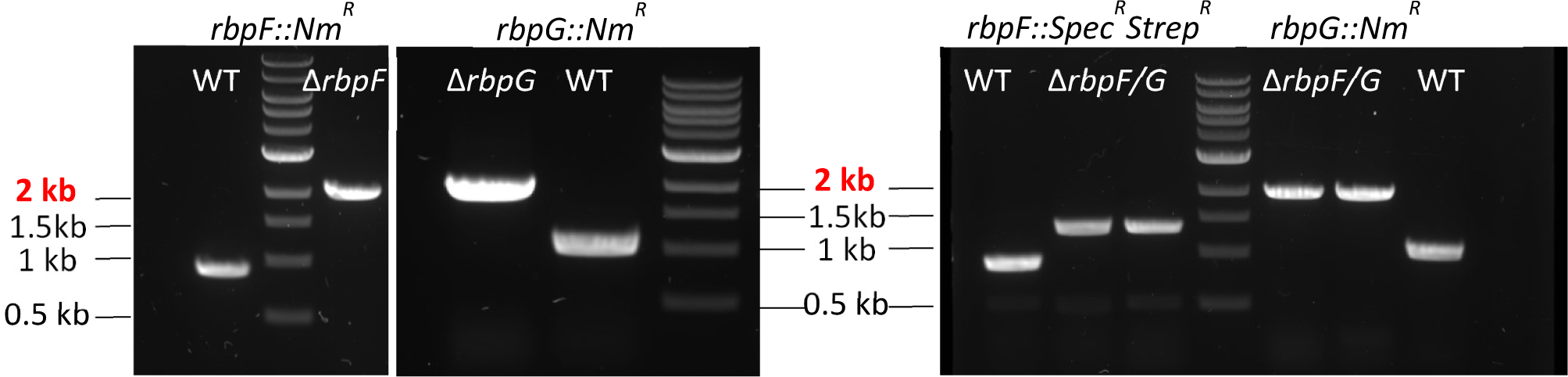
PCR segregation tests for *Anabaena* Δ*rbpF*, Δ*rbpG* and Δ*rbpF*/*G* mutants.

**FIG S4.**
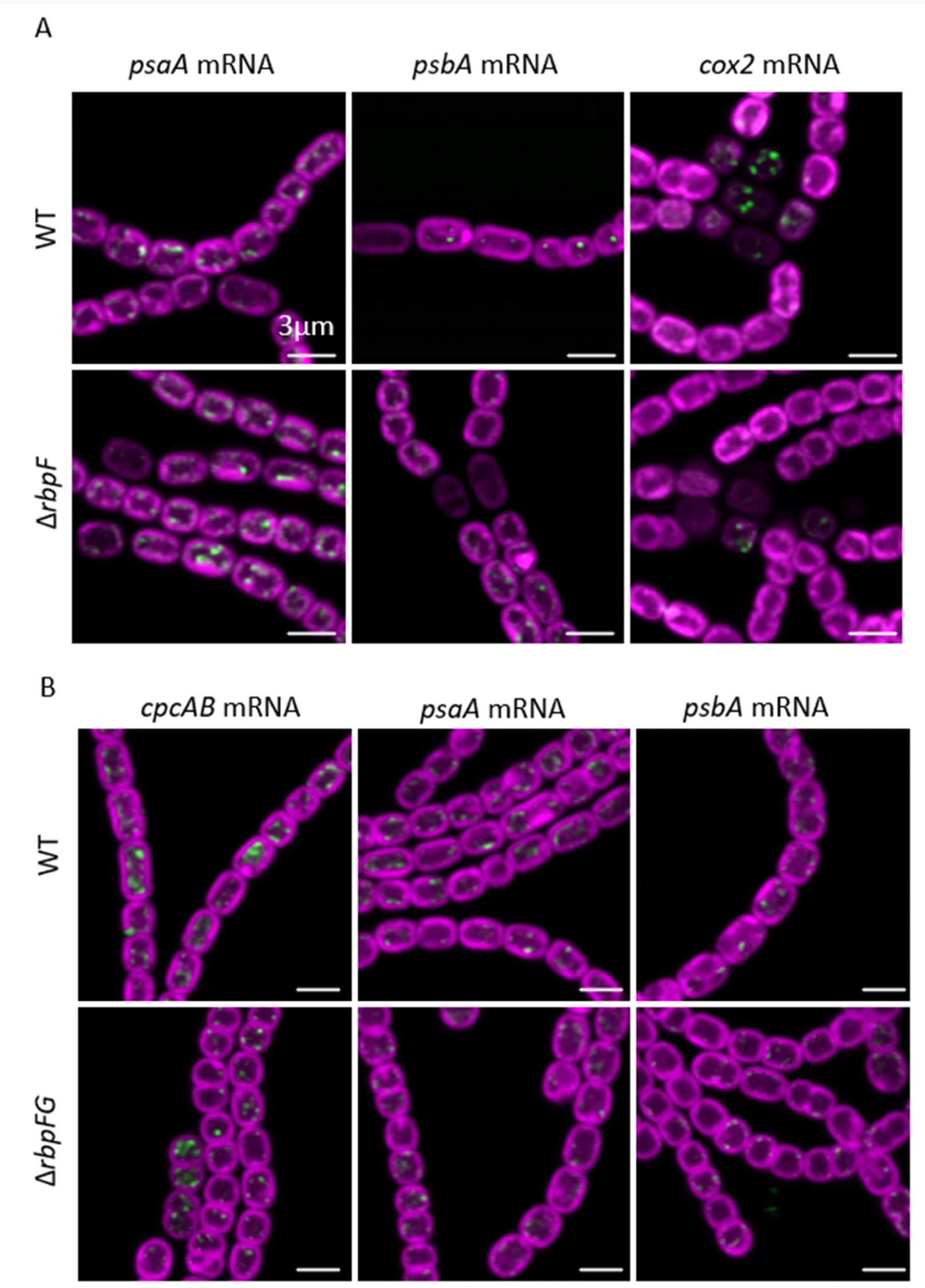
Location of mRNAs in *rbpF* and *rbpG* mutants.

**TABLE S1.**
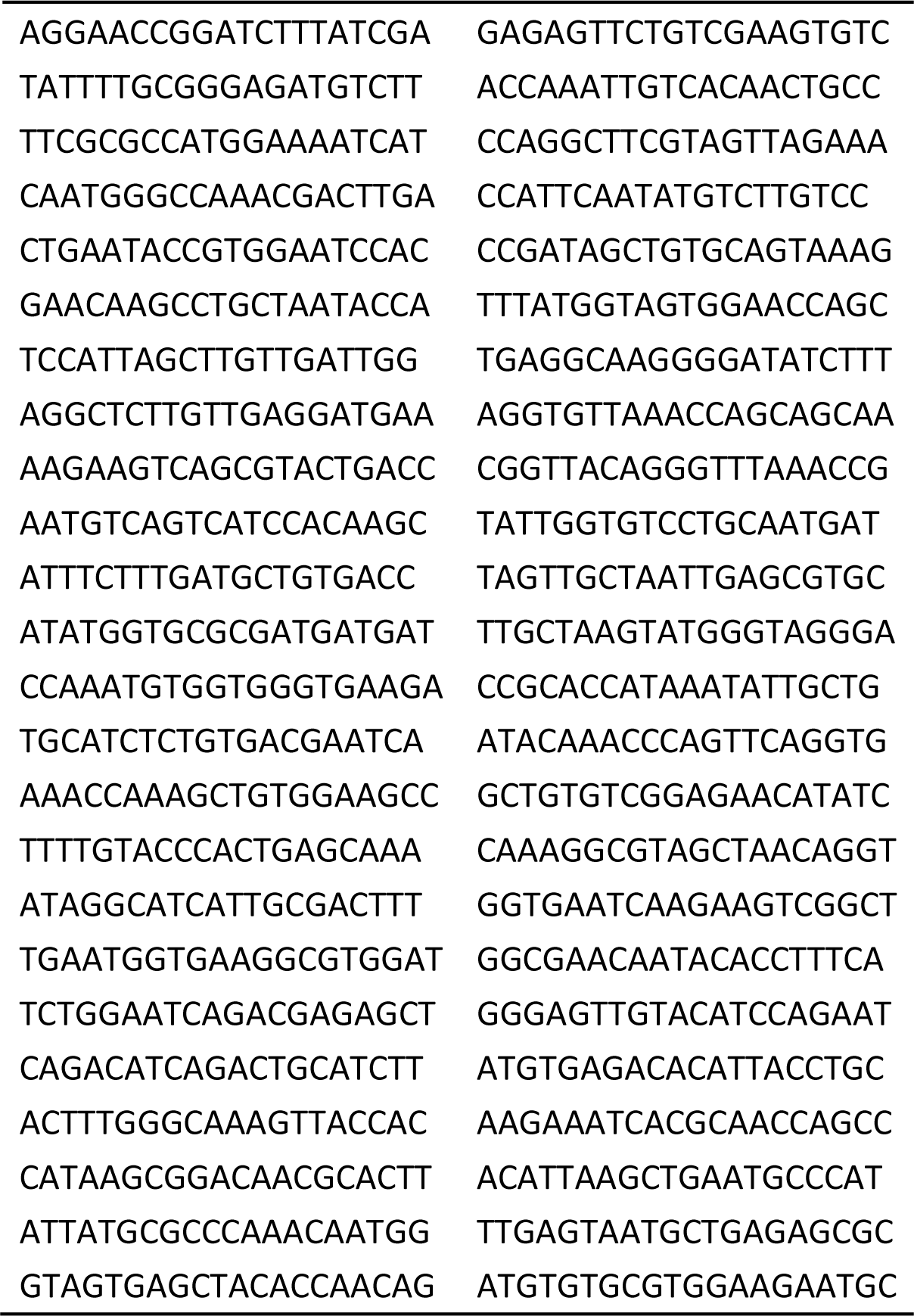
Set of 48 FISH probes for *Anabaena psaA* mRNA.

**TABLE S2.**
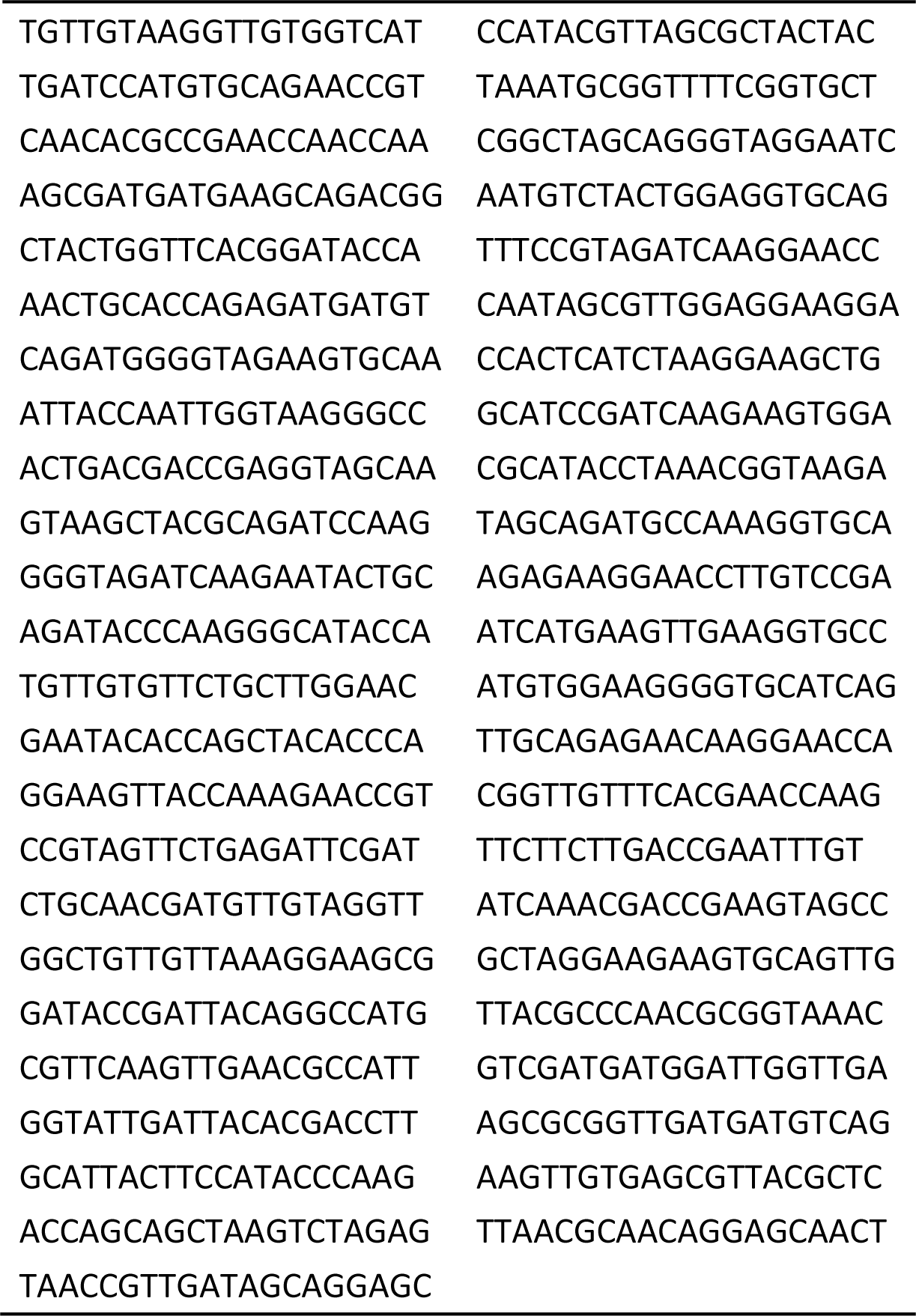
Set of 47 FISH probes for *Anabaena psbAI* mRNA.

**TABLE S3.**
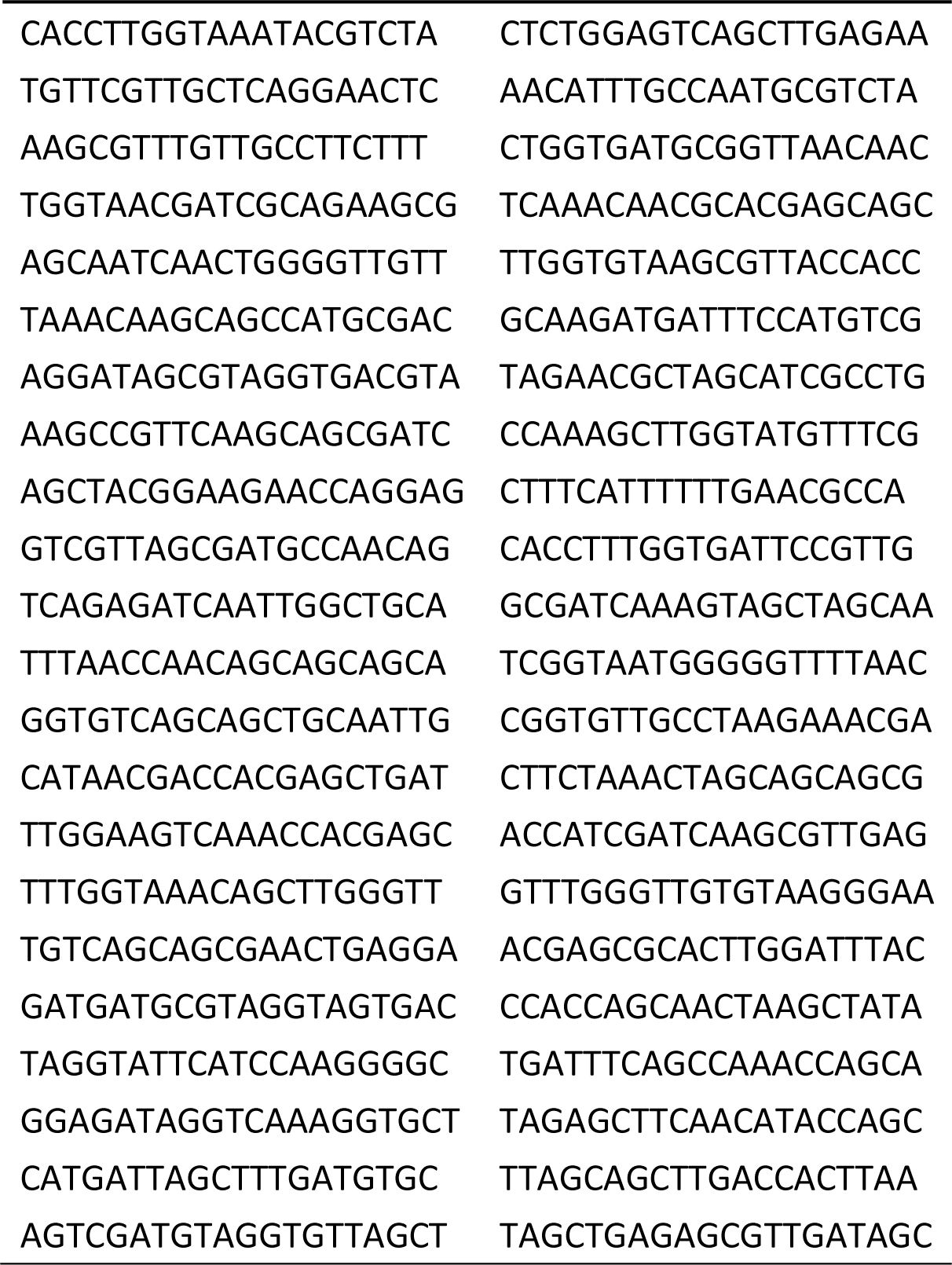
Set of 44 FISH probes for *Anabaena cpcAB* mRNA.

**TABLE S4.**
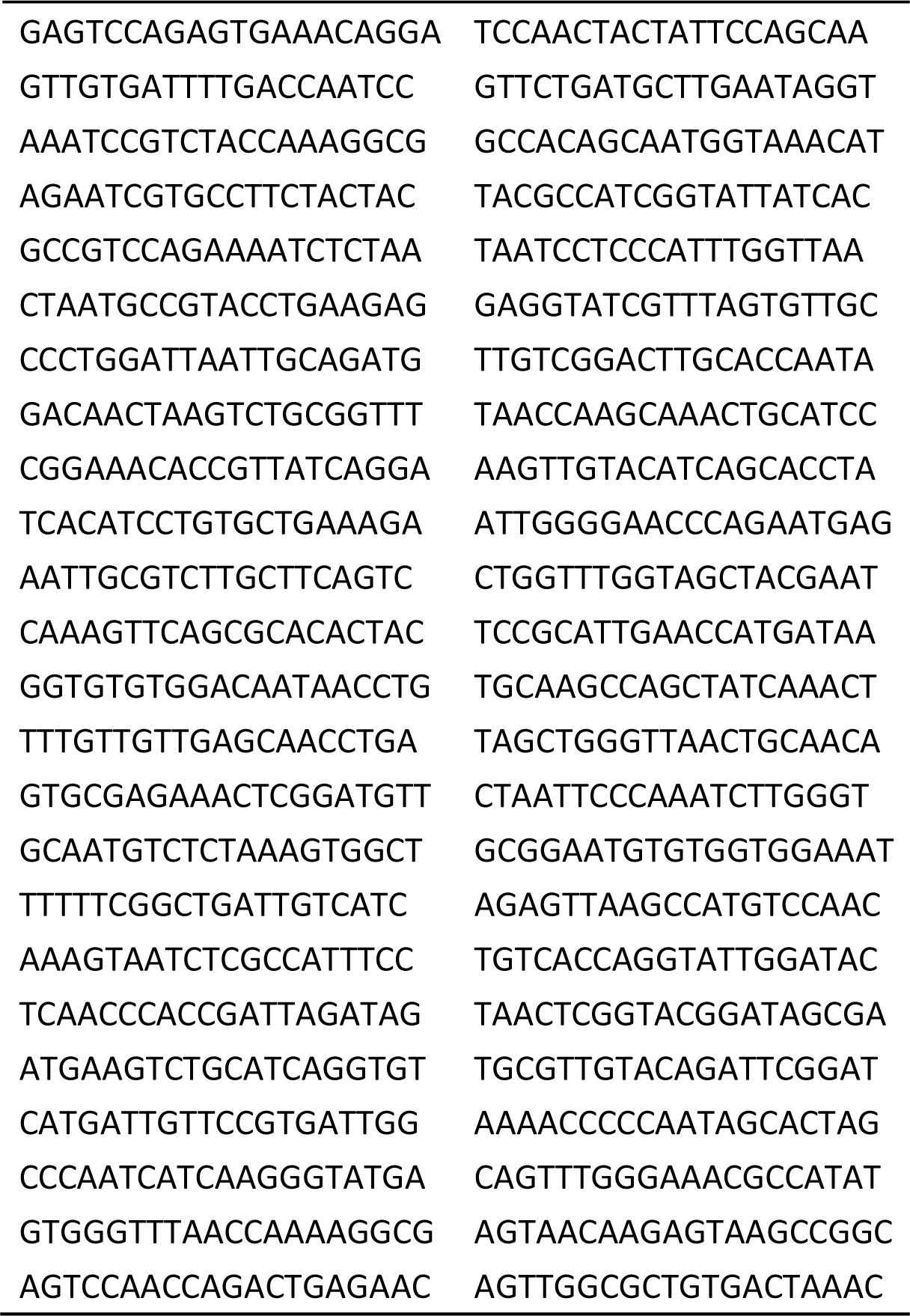
Set of 48 FISH probes for *Anabaena cox2* mRNA.

**TABLE S5.**
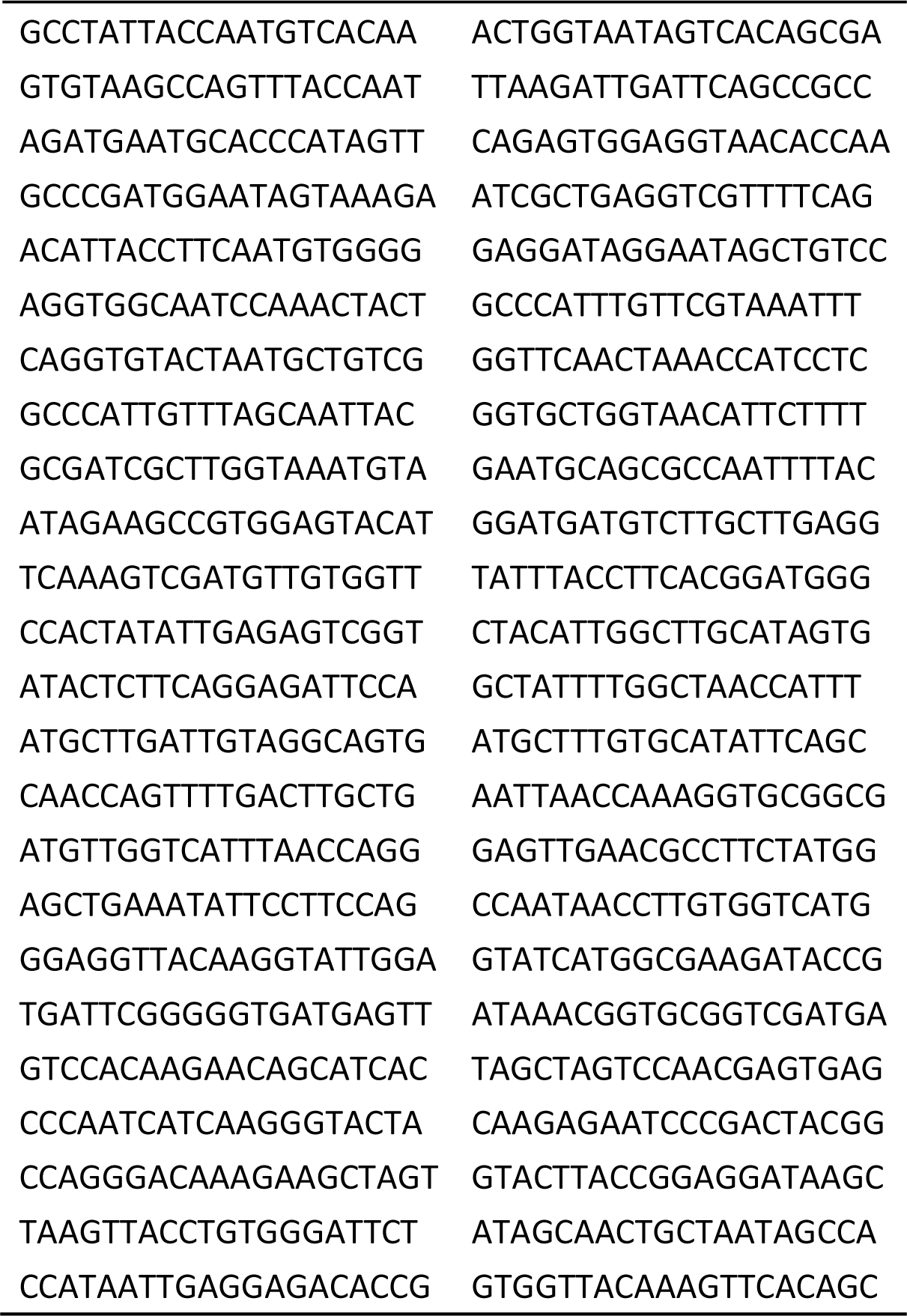
Set of 48 FISH probes for *Anabaena cox3* mRNA.

**TABLE S6.**
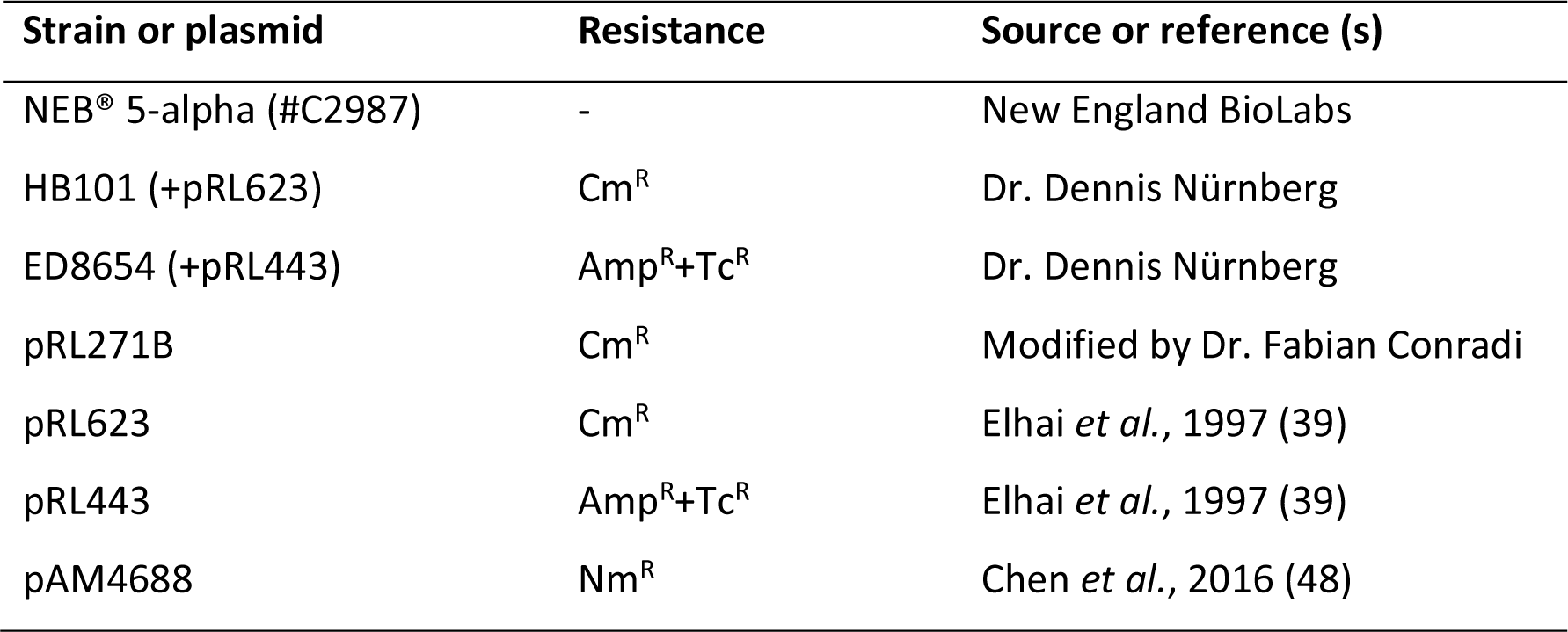
*E. coli* strains and plasmids used in this study.

| Strain or plasmid | Resistance | Source or reference (s) |
| --- | --- | --- |
| NEB® 5-alpha (#C2987) | - | New England BioLabs |
| HB101 (+pRL623) | Cm<sup>R</sup> | Dr. Dennis Nürnberg |
| ED8654 (+pRL443) | Amp<sup>R</sup>+Tc<sup>R</sup> | Dr. Dennis Nürnberg |
| pRL271B | Cm<sup>R</sup> | Modified by Dr. Fabian Conradi |
| pRL623 | Cm<sup>R</sup> | Elhai *et al.*, 1997 (39) |
| pRL443 | Amp<sup>R</sup>+Tc<sup>R</sup> | Elhai *et al.*, 1997 (39) |
| pAM4688 | Nm<sup>R</sup> | Chen *et al.*, 2016 (48) |

**TABLE S7.**
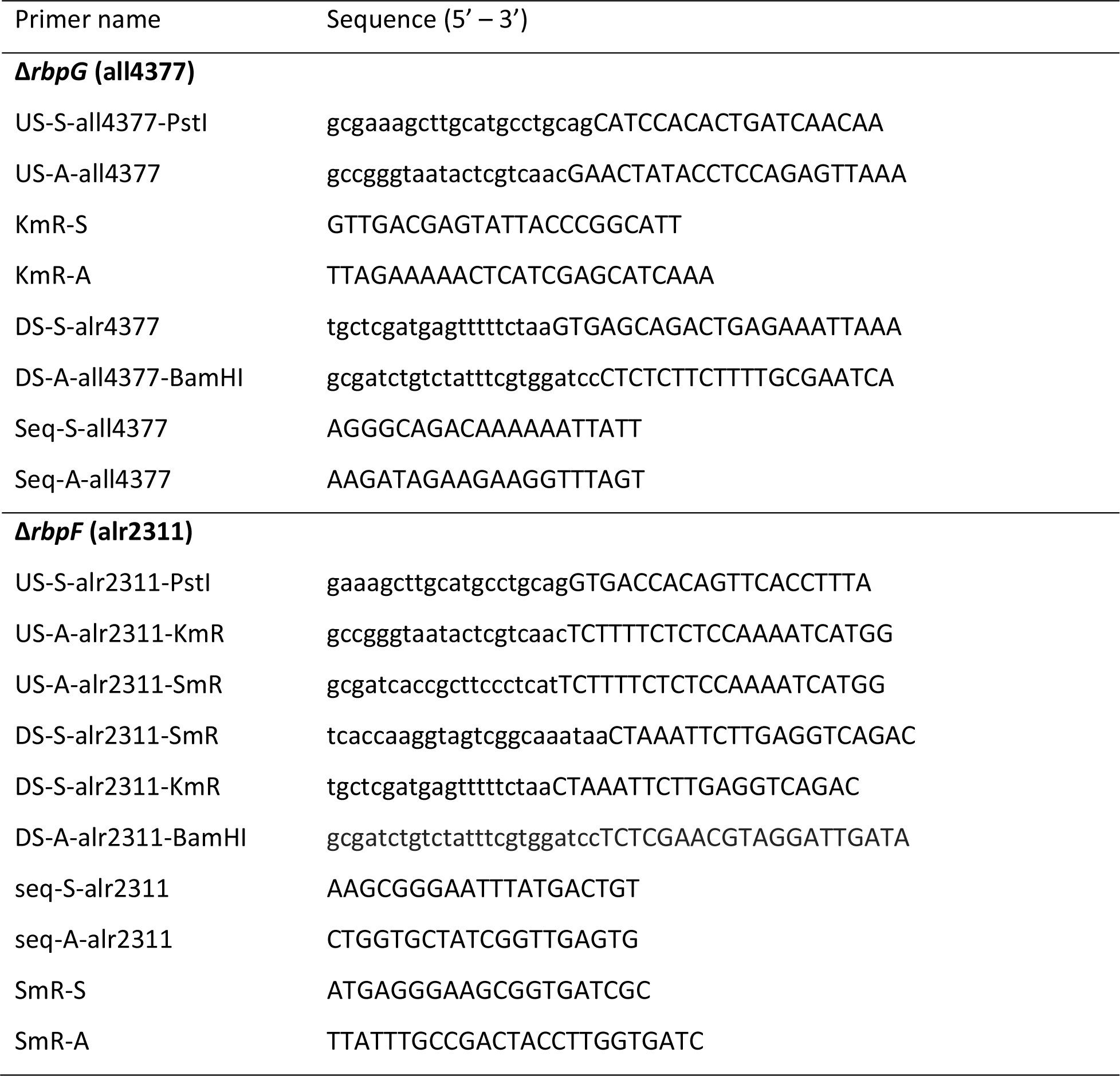
Oligonucleotide primers used to generate constructs for deletion of *Anabaena rbpG* and *rbpF*.

## REFERENCES

1. Nürnberg DJ, Mariscal V, Bornikoel J, Nieves-Morión M, Krauß N, Herrero A, Maldener I, Flores E, Mullineaux CW. 2015. Intercellular diffusion of a fluorescent sucrose analog via the septal Junctions in a filamentous cyanobacterium. MBio 6:1–12.

2. Harish, Seth K. 2020. Molecular circuit of heterocyst differentiation in cyanobacteria. J Basic Microbiol 60:738–745.

3. Zeng X, Zhang C-C. 2022. The making of a heterocyst in cyanobacteria. Annu Rev Microbiol 76:597–618.

4. Donze M, Haveman J, Schiereck P. 1972. Absence of Photosystem 2 in heterocysts of the blue-green alga *Anabaena*. Biochim Biophys Acta - Bioenerg 256:157–161.

5. Magnuson A, Cardona T. 2016. Thylakoid membrane function in heterocysts. Biochim Biophys Acta - Bioenerg 1857:309–319.

6. Santamaría-Gómez J, Mariscal V, Luque I. 2018. Mechanisms for protein redistribution in thylakoids of *Anabaena* during cell differentiation. Plant Cell Physiol 59:1860–1873.

7. Murry MA, Olafsen AG, Benemann JR. 1981. Oxidation of diaminobenzidine in the heterocysts of *Anabaena cylindrica*. Curr Microbiol 6:201–206.

8. Valladares A, Maldener I, Muro-Pastor AM, Flores E, Herrero A. 2007. Heterocyst development and diazotrophic metabolism in terminal respiratory oxidase mutants of the cyanobacterium *Anabaena* sp. strain PCC 7120. J Bacteriol 189:4425–4430.

9. Valladares A, Herrero A, Pils D, Schmetterer G, Flores E. 2003. Cytochrome *c* oxidase genes required for nitrogenase activity and diazotrophic growth in *Anabaena* sp. PCC 7120. Mol Microbiol 47:1239–1249.

10. Merino-Puerto V, Mariscal V, Mullineaux CW, Herrero A, Flores E. 2010. Fra proteins influencing filament integrity, diazotrophy and localization of septal protein SepJ in the heterocyst-forming cyanobacterium *Anabaena* sp. Mol Microbiol 75:1159–1170.

11. Merino-Puerto V, Mariscal V, Schwarz H, Maldener I, Mullineaux CW, Herrero A, Flores E. 2011. FraH is required for reorganization of intracellular membranes during heterocyst differentiation in *Anabaena* sp. strain PCC 7120. J Bacteriol 193:6815–6823.

12. Xu C, Wang B, Heng H, Huang J, Wan C. 2022. Comparative network biology discovers protein complexes that underline cellular differentiation in *Anabaena* sp. Mol Cell Proteomics 21:100224.

13. Corrales-Guerrero L, Mariscal V, Nürnberg DJ, Elhai J, Mullineaux CW, Flores E, Herrero A. 2014. Subcellular localization and clues for the function of the HetN factor influencing heterocyst distribution in *Anabaena* sp. strain PCC 7120. J Bacteriol 196:3452–60.

14. Santamaría-Gómez J, Ochoa de Alda JAG, Olmedo-Verd E, Bru-Martínez R, Luque I. 2016. Sub-cellular localization and complex formation by aminoacyl-tRNA synthetases in cyanobacteria: evidence for interaction of membrane-anchored ValRS with ATP synthase. Front Microbiol 7:857.

15. Olmedo-Verd E, Santamaría-Gómez J, Ochoa de Alda JAG, de Pouplana LR, Luque I. 2011. Membrane anchoring of aminoacyl-tRNA synthetases by convergent acquisition of a novel protein domain. J Biol Chem 286:41057–41068.

16. Mahbub M, Hemm L, Yang Y, Kaur R, Carmen H, Engl C, Huokko T, Riediger M, Watanabe S, Liu L-N, Wilde A, Hess WR, Mullineaux CW. 2020. mRNA localization, reaction centre biogenesis and thylakoid membrane targeting in cyanobacteria. Nat Plants 6:1179–1191.

17. Skinner SO, Sepúlveda LA, Xu H, Golding I. 2013. Measuring mRNA copy number in individual *Escherichia coli* cells using single-molecule fluorescent *in situ* hybridization. Nat Protoc 8:1100–1113.

18. Kaneko T, Nakamura Y, Wolk CP, Kuritz T, Sasamoto S, Watanabe A, Iriguchi M, Ishikawa A, Kawashima K, Kimura T, Kishida Y, Kohara M, Matsumoto M, Matsuno A, Muraki A, Nakazaki N, Shimpo S, Sugimoto M, Takazawa M, Yamada M, Yasuda M, Tabata S. 2001. Complete genomic sequence of the filamentous nitrogen-fixing cyanobacterium *Anabaena* sp. strain PCC 7120. DNA Res 8:205–213.

19. Sicora CI, Appleton SE, Brown CM, Chung J, Chandler J, Cockshutt AM, Vass I, Campbell DA. 2006. Cyanobacterial psbA families in *Anabaena* and *Synechocystis* encode trace, constitutive and UVB-induced D1 isoforms. Biochim Biophys Acta - Bioenerg 1757:47–56.

20. Salem K, van Waasbergen LG. 2004. Photosynthetic electron transport controls expression of the High Light Inducible gene in the cyanobacterium *Synechococcus elongatus* strain PCC 7942. Plant Cell Physiol 45:651–658.

21. Golden JW, Yoon H-S. 1998. Heterocyst formation in *Anabaena*. Curr Opin Microbiol 1:623–629.

22. Pernil R, Schleiff E. 2019. Metalloproteins in the biology of heterocysts. Life.

23. Ehira S, Hamano T, Hayashida T, Kojima K, Nakamoto H, Hiyama T, Ohmori M, Shivaji S, Sato N. 2003. Conserved temperature-dependent expression of RNA-binding proteins in cyanobacteria with different temperature optima. FEMS Microbiol Lett 225:137–142.

24. Maruyama K, Sato N, Ohta N. 1999. Conservation of structure and cold-regulation of RNA-binding proteins in cyanobacteria: Probable convergent evolution with eukaryotic glycine-rich RNA-binding proteins. Nucleic Acids Res 27:2029–2036.

25. Hamano T, Murakami S, Takayama K, Ehira S, Maruyama K, Kawakami H, Morita EH, Hayashi H, Sato N. 2004. Characterization of RNA-binding properties of three types of RNA-binding proteins in *Anabaena* sp. PPC 7120. Cell Mol Biol (Noisy-le-grand) 50:613–624.

26. Zhang Y, Wu D, Wang Y, Xu X. 2022. Two types of C-terminal regions of RNA-binding proteins play distinct roles in stress tolerance of *Synechocystis* sp. PCC 6803. FEMS Microbiol Lett 369.

27. Komenda J, Sobotka R, Nixon PJ. 2012. Assembling and maintaining the Photosystem II complex in chloroplasts and cyanobacteria. Curr Opin Plant Biol 15:245–251.

28. Graan T, Ort DR. 1986. Detection of oxygen-evolving Photosystem II centers inactive in plastoquinone reduction. Biochim Biophys Acta - Bioenerg 852:320–330.

29. Mitschke J, Vioque A, Haas F, Hess WR, Muro-Pastor AM. 2011. Dynamics of transcriptional start site selection during nitrogen stress-induced cell differentiation in *Anabaena* sp. PCC7120. Proc Natl Acad Sci 108:20130–20135.

30. Kushige H, Kugenuma H, Matsuoka M, Ehira S, Ohmori M, Iwasaki H. 2013. Genome-wide and heterocyst-specific circadian gene expression in the filamentous cyanobacterium *Anabaena* sp. strain PCC 7120. J Bacteriol 195:1276–1284.

31. Aldea MR, Mella-Herrera RA, Golden JW. 2007. Sigma factor genes *sigC*, *sigE*, and *sigG* are upregulated in heterocysts of the cyanobacterium *Anabaena* sp. strain PCC 7120. J Bacteriol 189:8392–8396.

32. Rajagopalan R, Callahan SM. 2010. Temporal and spatial regulation of the four transcription start sites of hetR from *Anabaena* sp. strain PCC 7120. J Bacteriol 192:1088– 1096.

33. Constant S, Perewoska I, Alfonso M, Kirilovsky D. 1997. Expression of the *psbA* gene during photoinhibition and recovery in *Synechocystis* PCC 6714: Inhibition and damage of transcriptional and translational machinery prevent the restoration of photosystem II activity. Plant Mol Biol 34:1–13.

34. Herranen M, Aro E-M, Tyystjärvi T. 2001. Two distinct mechanisms regulate the transcription of photosystem II genes in Synechocystis sp. PCC 6803. Physiol Plant 112:531–539.

35. Nevo-Dinur K, Nussbaum-Shochat A, Ben-Yehuda S, Amster-Choder O. 2011. Translation-independent localization of mRNA in *E. coli*. Science 331:1081–4.

36. Mahbub M, Mullineaux CW. 2023. Locations of membrane protein production in a cyanobacterium. J Bacteriol in revision.

37. Heinz S, Rast A, Shao L, Gutu A, Gügel IL, Heyno E, Labs M, Rengstl B, Viola S, Nowaczyk MM, Leister D, Nickelsen J. 2016. Thylakoid membrane architecture in *Synechocystis* depends on CurT, a homolog of the granal CURVATURE THYLAKOID1 proteins. Plant Cell 28:2238–2260.

38. Rippka R, Deruelles J, Waterbury JB, Herdman M, Stanier RY. 1979. Generic assignments, strain histories and properties of pure cultures of cyanobacteria. J Gen Microbiol 111:1– 61.

39. Elhai J, Vepritskiy A, Muro-Pastor AM, Flores E, Wolk CP. 1997. Reduction of conjugal transfer efficiency by three restriction activities of *Anabaena* sp. strain PCC 7120. J Bacteriol 179:1998–2005.

40. Elhai J, Wolk CP. 1988. Conjugal transfer of DNA to cyanobacteria, p. 747–754. In Packer, L, Glazer, AN (eds.), Methods in Enzymology Vol.167: Cyanobacteria. Academic Press, San Diego.

41. Schindelin J, Arganda-Carreras I, Frise E, Kaynig V, Longair M, Pietzsch T, Preibisch S, Rueden C, Saalfeld S, Schmid B, Tinevez J-Y, White DJ, Hartenstein V, Eliceiri K, Tomancak P, Cardona A. 2012. Fiji: an open-source platform for biological-image analysis. Nat Methods 9:676–682.

42. Myers J, Graham J-R, Wang RT. 1980. Light harvesting in *Anacystis nidulans* studied in pigment mutants. Plant Physiol 66:1144–1149.

43. Thompson JD, Higgins DG, Gibson TJ. 1994. CLUSTAL W: improving the sensitivity of progressive multiple sequence alignment through sequence weighting, position-specific gap penalties and weight matrix choice. Nucleic Acids Res 22:4673–4680.

44. Hall TA. 1999. BioEdit: a user-friendly biological sequence alignment editor and analysis program for Windows 95/98/NT. Nucleic Acids Symp Ser 41:95–98.

45. Rzhetsky A, Nei M. 1992. A Simple Method for Estimating and Testing Minimum-Evolution Trees. Mol Biol Evol 9:945.

46. Kumar S, Stecher G, Li M, Knyaz C, Tamura K. 2018. MEGA X: Molecular Evolutionary Genetics Analysis across Computing Platforms. Mol Biol Evol 35:1547–1549.

47. Zuckerkandl E, Pauling L. 1965. Evolutionary Divergence and Convergence in Proteins, p. 97–166. In Bryson, V, Vogel, H (eds.), Evolving Genes and Proteins. Academic Press.

48. Chen Y, Taton A, Go M, London RE, Pieper LM, Golden SS, Golden JW. 2016. Self-replicating shuttle vectors based on pANS, a small endogenous plasmid of the unicellular cyanobacterium *Synechococcus elongatus* PCC 7942. Microbiol (United Kingdom) 162:2029–2041.

